# Transcriptional regulation of photoprotection in dark-to-light transition - more than just a matter of excess light energy

**DOI:** 10.1101/2021.10.23.463292

**Authors:** Petra Redekop, Emanuel Sanz-Luque, Yizhong Yuan, Gaelle Villain, Dimitris Petroutsos, Arthur R. Grossman

## Abstract

In nature, photosynthetic organisms are exposed to different light spectra and intensities depending on the time of day and atmospheric and environmental conditions. When photosynthetic cells absorb excess light, they induce non-photochemical quenching to avoid photo-damage and trigger expression of ‘photoprotective’ genes. In this work, we used the green alga *Chlamydomonas reinhardtii* to assess the impact of light intensity, light quality, wavelength, photosynthetic electron transport and CO_2_ on induction of the ‘photoprotective’ genes (*LHCSR1*, *LHCSR3* and *PSBS*) during dark-to-light transitions. Induction (mRNA accumulation) occurred at very low light intensity, was independently modulated by blue and UV-B radiation through specific photoreceptors, and only *LHCSR3* was strongly controlled by CO_2_ levels through a putative enhancer function of CIA5, a transcription factor that controls genes of the carbon concentrating mechanism. We propose a model that integrates inputs of independent signaling pathways and how they may help the cells anticipate diel conditions and survive in a dynamic light environment.

## INTRODUCTION

Light absorption and its conversion into chemical energy by photosynthetic organisms is an essential process for almost all life on our planet. Photosynthetic organisms have evolved to efficiently capture light energy when the intensity is low, and quench absorbed excitation energy when it exceeds the photon flux density needed to saturate photosynthetic electron transport. Excess light leads to the generation of reactive oxygen species (ROS) that cause cellular damage and even cell death. Photoprotection requires the activities of a set of proteins that functions to dissipate excess absorbed light energy before it is used to drive reaction center function. In the green alga *Chlamydomonas reinhardtii* (Chlamydomonas throughout), LHCSR3 (encoded by *LHCSR3.1* and *LHCSR3.2*, that only differ slightly in their promoters), LHCSR1 and PSBS (encoded by *PSBS1* and *PSBS2*; the proteins differ by one amino acid) are often described as photoprotective proteins that accumulate in response to high light (HL) and UV-B radiation (280-315 nm) (Allorent et al., 2016; Ballottari et al., 2016; Bonente et al., 2011; Correa-Galvis et al., 2016; Dinc et al., 2016; Peers et al., 2009; Tibiletti et al., 2016). These photoprotective proteins are critical for rapid non-photochemical quenching (NPQ) of excess absorbed light energy through a process designated qE (energy dependent quenching). For the Chlamydomonas *PSBS* genes, the transcript and proteins accumulate transiently in response to HL and UV-B radiation (Allorent et al., 2016; Correa-Galvis et al., 2016; Tibiletti et al., 2016). The exact function of PSBS in Chlamydomonas needs further elucidation, although it was found to positively impact acclimation to HL (Redekop et al., 2020) and studies in vascular plants have demonstrated that it functions in conjunction with the xanthophyll cycle and a ΔpH across the thylakoid membranes to elicit qE (Bonente et al., 2008; Peers et al., 2009; Sacharz et al., 2017). LHCSR proteins (LHCSX in diatoms) are the dominant ‘quenching’ proteins in algae and while present in moss, no orthologs have been identified in vascular plants (Alboresi et al., 2010; Bailleul et al., 2010; Pinnola, 2019; Teramoto et al., 2002).

To elicit an efficient photoprotection response, cells would need to accumulate the photoprotective proteins before or very soon after exposure to HL. At the transcriptional level, *LHCSR* and *PSBS* genes are strongly induced during the dark to light transitions, especially when this transition is abrupt (Strenkert et al., 2019). However, the signals that prime cells for eliciting photoprotective processes are still not well understood. There are many questions concerning the mechanisms and the factors controlling accumulation of *LHCSR3*, *LHCSR1* and *PSBS* transcripts and the encoded proteins. Expression of these genes is impacted by specific photoreceptors including the UV RESISTANCE LOCUS 8 (UVR8) (Allorent et al., 2016; Tilbrook et al., 2016) and the blue light photoreceptor phototropin (PHOT) (Petroutsos et al., 2016).

UVR8 in *Arabidopsis thaliana* is homodimeric and absorbs UV-B radiation through tryptophan residues with a peak in its action spectrum at 260-280 nm (Jiang et al., 2012). The absorption of UV-B radiation by UVR8 causes monomerization of the photoreceptor and facilitates its interactions with CONSTITUTIVELY PHOTOMORPHOGENIC 1 (COP1), a protein with E3 ubiquitin ligase activity (Hoecker, 2017; X. Huang et al., 2014), and SUPPRESSOR OF PHYA-105 1 (SPA1) (Menon et al., 2016). This complex prevents degradation of ELONGATED HYPOCOTYL 5 (HY5), a transcription factor that activates gene expression in response to UV-B radiation (Heijde & Ulm, 2012), including the responses associated with acclimation of plants to excess absorbed excitation energy (Favory et al., 2009).

Chlamydomonas UVR8, COP1 and SPA1 orthologs also have functions related to quenching excess absorbed light energy (Tilbrook et al., 2016; Gabilly et al., 2019). COP1 and SPA1, along with CULLIN4 (CUL4) and DAMAGED DNA BINDING 1 (DDB1), form an E3 ubiquitin ligase complex. This complex controls light dependent transcription of Chlamydomonas *LHCSR* and *PSBS* genes and appears to be critical for suppressing expression of the genes associated with qE in LL grown cells (Gabilly et al., 2019), which involves ubiquitination of a complex formed by two transcription factors, CrCO and NF-Ys. These factors are required for eliciting maximum induction of proteins associated with qE (Tokutsu, Fujimura-Kamada, Yamasaki, et al., 2019). The absorption of UV-B radiation by UVR8 leads to the interaction of monomeric UVR8 with COP1, which promotes SPA1/COP1 dissociation from Cr-CO, which in turn blocks Cr-CO ubiquitination and degradation, and allows the formation of a stable Cr-CO/NF-Ys transcription complex that elicits increased accumulation of *LHCSR* and *PSBS* transcripts (Tokutsu et al., 2021; Tokutsu, et al., 2019).

The blue-light photoreceptor PHOT1 consists of two similar photosensory LOV1/2 (Light, Oxygen and Voltage sensitive) domains at the N terminus and a serine/ threonine kinase domain at the C terminus. Upon perception of blue light by LOV1/2 the photoreceptor is autophosphorylated and activates cellular responses that promote plant growth under weak light conditions. Higher plants code for two PHOTs, PHOT1 and PHOT2, with distinct and overlapping functions including phototropism, stomatal opening, chloroplast movement, and cotyledon and leaf expansion (Christie, 2007). In Chlamydomonas, there is a single PHOT (designated PHOT1) that controls expression of genes for enzymes in the chlorophyll and carotenoid biosynthesis pathways (Im et al., 2006), regulates multiple steps of the sexual life cycle (K. Huang & Beck, 2003) and acts as light regulator of phototaxis (Trippens et al., 2012). A link between PHOT1 and activation of the *LHCSR3* gene has also been established (Petroutsos et al., 2016).

Recent evidence suggests that *LHCSR1*, *LHCSR3* and *PSBS* in Chlamydomonas are transcribed in response to HL and UV-B radiation, with *LHCSR3* specifically requiring a PHOT-dependent and a light-dependent signal generated in the chloroplast that is still to be defined (Petroutsos et al., 2016). *LHCSR1* (Tibiletti et al., 2016) and *PSBS* (Allorent & Petroutsos, 2017) were previously proposed to be regulated by high white light by an unknown pathway. The work presented in this manuscript provides new insights into the features of radiation that impact expression of these three photoprotective genes. We found that a strong induction of all three genes is observed following a shift from dark to LL; this LL-elicited increase in transcript abundances is mostly independent of photosynthetic electron transport (PET) but dependent on PHOT1. We also demonstrate that UV-B radiation independent of photosynthetically active radiation (PAR) can cause maximum accumulation of *LHCSR1*, *PSBS*, and near maximum accumulation of *LHCSR3* transcripts; this response is mostly suppressed in the *uvr8* mutant. Furthermore, *LHCSR3* is strongly regulated by CIA5 through a potential enhancer function that is needed to elevate expression under all conditions, while there is little or no CIA5-dependent control of *LHCSR1* or *PSBS* at the transcript level, although *PSBS* may be impacted to a minor extend by high CO_2_. Additionally, we discuss the potential integration of these signals in nature.

## RESULTS

### Photoprotective genes are induced at low light intensities

Light intensity and quality, including levels of UV-B, markedly change over the diel cycle. An hourly characterization of PAR and UV-B intensities were tracked from dawn to dusk in July in California under generally sunny skies, with some cloud cover at 9:00-10:00 AM. Both PAR and UV-B intensities gradually increased, with a broad PAR peak reaching maximal values between 11:00 AM and 4:00 PM and a narrower UV-B peak (**Figure 1A**). As shown in **Figure 1A** (note arrow), clouds impact the intensity of PAR much more than that of UV-B radiation. Furthermore, from 6:00 AM to 8:00 AM the PAR intensity increased steeply from less than 100 µmol photons m^-2^ s^-1^ to more than 1000 µmol photons m^-2^ s^-1^ and then more gradually, reaching a peak of ∼2000 µmol photons m^-2^ s^-1^ that is sustained over a period of 5-6 h. Hence, acclimation to HL occurs progressively, starting under very low light conditions in the early morning, with rapidly increasing intensities over the course of ∼4 h, reaching intensities that result in the hyper-saturation of photosynthesis (photosynthesis saturates at ∼400-1000 µmol photons m^-2^ s^-1^) that is sustained over a large proportion of the day (Formighieri et al., 2012; Polle et al., 2000).

**Figure 1:**
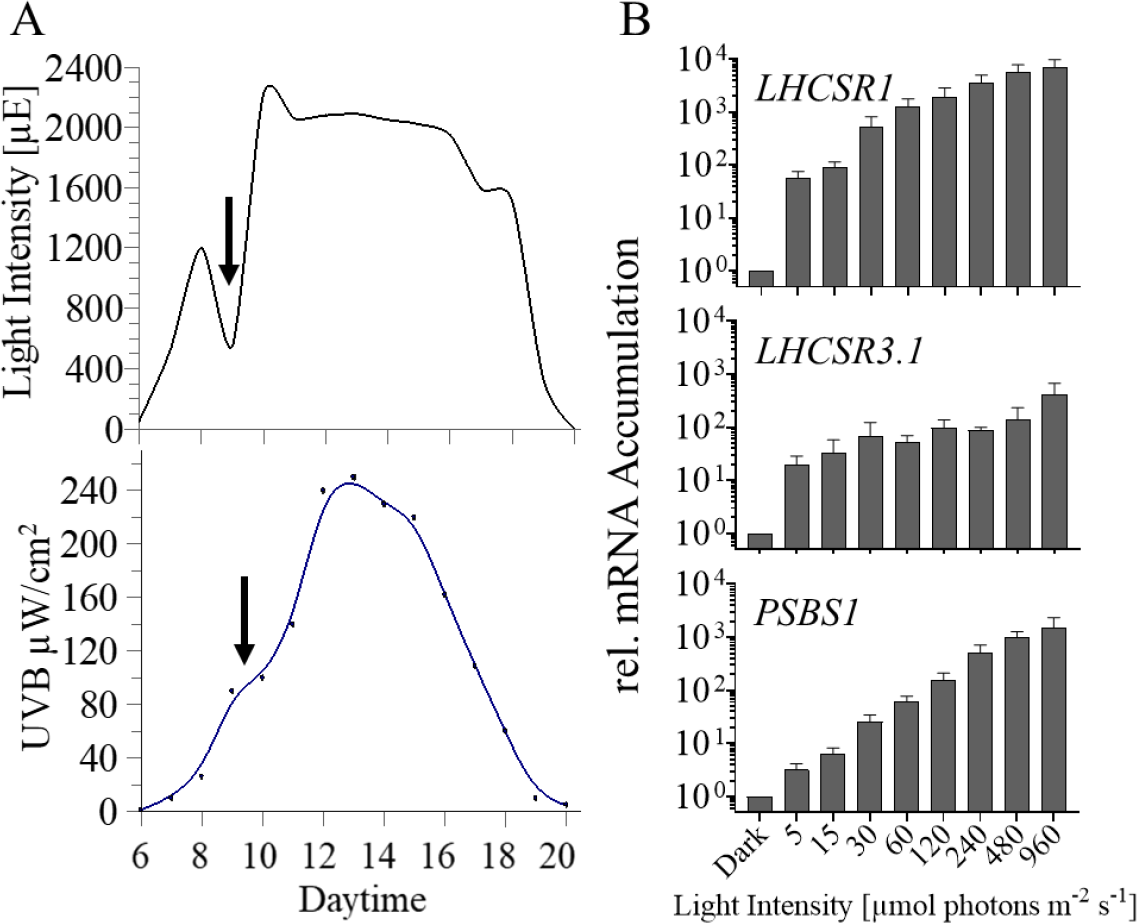
PAR and UV-B intensity from sunrise to sunset and changes in *LHCSRs* and *PSBS1* transcript levels after one hour of irradiation at various light intensities. **A)** The intensity of PAR and UV-B were monitored from dawn to dusk (July, California). **B**) Accumulation of *LHCSR1*, *LHCSR3.1* and *PSBS1* transcripts following incubation for 1 h at different light intensities. WT CC125 cells were grown in TAP at LL (30 µmol photons m^-2^ s^-1^) and then transferred to the dark for 24 h (in TAP, maintaining a fixed carbon source in the dark). Cells were then transferred to HSM (photoautotrophic conditions) and maintained for 2 additional h in the dark (to reduce intracellular levels of acetate and bring cells into a physiologically relevant state in which they would most effectively quench excessive absorbed light) prior to a 1 h light exposure at each of the indicated intensities. Abundances of each of the three transcripts were normalized to their dark control value. n=3+SD. Statistical analyses and P-values are listed in **Supplementary Table 1B**.

We postulated that algae accumulate transcripts from photoprotective genes (often described as ‘high light responsive’ genes) even in the early morning when there is a dark to light transition (when the light intensities are sub-saturating). While the amount of photoprotective proteins like LHCSR3 are stable in the dark under laboratory conditions (Nawrocki et al., 2019), this may not occur in the natural environment where the cells often experience dynamic and extreme conditions (e.g., nutrient limitation, anoxia, etc.) that might lead to protein turnover. Hence, the rapid elevations of transcripts encoding the photoprotective proteins during a dark to light transition might serve to prime the cells for a marked increase in radiation that occurs during the first few hours of light in the morning. Levels of transcripts from *LHCSR1*, *LHCSR3.1* and *PSBS1* genes were analyzed in dark-acclimated wild type (WT) cells (the WT is CC125 unless otherwise stated) after exposure to 1 h of PAR at 5, 15, 30, 60, 120, 240, 480 and 960 µmol photons m^-2^ s^-1^. The *LHCSR1* and *LHCSR3.1* genes showed strong induction even when exposed to very low light. There was a 56-fold increase for *LHCSR1* mRNA and a 20-fold increase for *LHCSR3.1* mRNA at 5 µmol photons m^-2^ s^-1^ compared to cells maintained in the dark. Furthermore, although maximum levels of mRNA accumulation from these two genes were observed at the highest light intensities (960 µmol photons m^-2^ s^-1^), it is remarkable that *LHCSR1* transcript levels showed a 500-fold induction after exposure to only 30 µmol photons m^-2^ s^-1^, and *LHCSR3.1* exhibited an induction of >60-fold at 30 µmol photons m^-2^ s^-1^. On the other hand, *PSBS1* was the least sensitive to low intensity radiation (e.g., 3-fold at 5 µmol photons m^-2^ s^-1^), and displayed a gradual, continuous increase in the level of its mRNA with increasing light intensity (**Figure 1B**, **Supplementary Figure 1**). The *LHCSR1* transcript also exhibited a gradual, continuous increase in transcript accumulation between 5 and 960 µmol photons m^-2^ s^-1^ (∼2 orders of magnitude) (**Figure 1B**, **Supplementary Figure 1**). Of the three transcripts, *LHCSR3.1* exhibited the lowest additional increase in transcript accumulation following its initial sharp rise at 5 µmol photons m^-2^ s^-1^; the difference between transcript abundance at 5 and 960 µmol photons m^-2^ s^-1^ is ∼20 fold. This increase was gradual from 5 to 480 µmol photons m^-2^ s^-1^, with a further increase of ∼2X between 480 and 960 µmol photons m^-2^ s^-1^. Additionally, we compared the levels of transcript accumulation across a light intensity gradient for the three genes from two different WT strains, CC125 and CC124 (**Supplementary Figure 1A**), which demonstrated that, although similar patterns were observed, there were differences in the sensitivity of the two strains to light intensity, most likely the consequence of genetic differences between them (Gallaher et al., 2015).

### *LHCSR1*, *LHCSR3*.*1* and *PSBS1* transcripts accumulate even when photosynthesis is blocked

In previous works, it was concluded that the maximum accumulation of the LHCSR3 protein required active linear electron transport but was independent of the redox state of the PQ pool. This conclusion was supported by the findings that accumulation of the LHCSR3 protein was inhibited in the presence of either DCMU [blocks Q_b_ binding site of photosystem (PS) II; the PQ pool becomes oxidized] or DBMIB (blocks electron transfer at the Qo site of the cytochrome *b*_6_ *f* complex; the PQ pool becomes reduced) and also in mutants devoid of either PSII (ΔPSII) or PSI (ΔPSI) (Petroutsos, Busch, Janssen, et al., 2011; Roberts & Kramer, 2001; Trebst, 2007). The lack of LHCSR3 protein accumulation in DCMU-treated samples correlated with essentially no increase of the mRNA from *LHCSR3.1* and *LHCSR3.2* in cells transferred from LL to HL in the presence of DCMU (Maruyama et al., 2014; Petroutsos et al., 2016). This previous work suggested a linear PET requirement for *LHCSR3* induction.

We examined the impact of DCMU on accumulation of the *LHCSR1*, *LHCSR3.1* and *PSBS1* transcripts following a dark-to-light transition. Congruent with the data in **Figure 1**, **Figure 2** shows that a transition from dark to LL (30 µmol photons m^-2^ s^-1^ for 1 h) strongly impacts *LHCSR1* and *LHCSR3.1* transcript levels, with less of an impact on the *PSBS1* transcript level. This low light exposure strongly induces the three genes even in the presence of DCMU (**Figure 2**). The *LHCSR3.1* transcript abundance increased 118-fold and 551-fold, in LL and HL respectively, and DCMU suppressed these increases by ∼50 %, indicating that, in the absence of linear electron flow, the cells can still induce expression of this gene by ∼55-fold and 236-fold in LL and HL, respectively. For *LHCSR1*, DCMU caused a similar reduction in transcript accumulation in LL but not in HL. The absence of a significant DCMU-mediated effect on *LHCSR1* gene expression under HL agrees with previously published data (Maruyama et al., 2014). There was no effect of DCMU on *PSBS1* mRNA accumulation in either LL or HL, suggesting that *PSBS1* expression is completely independent of linear electron flow.

**Figure 2:**
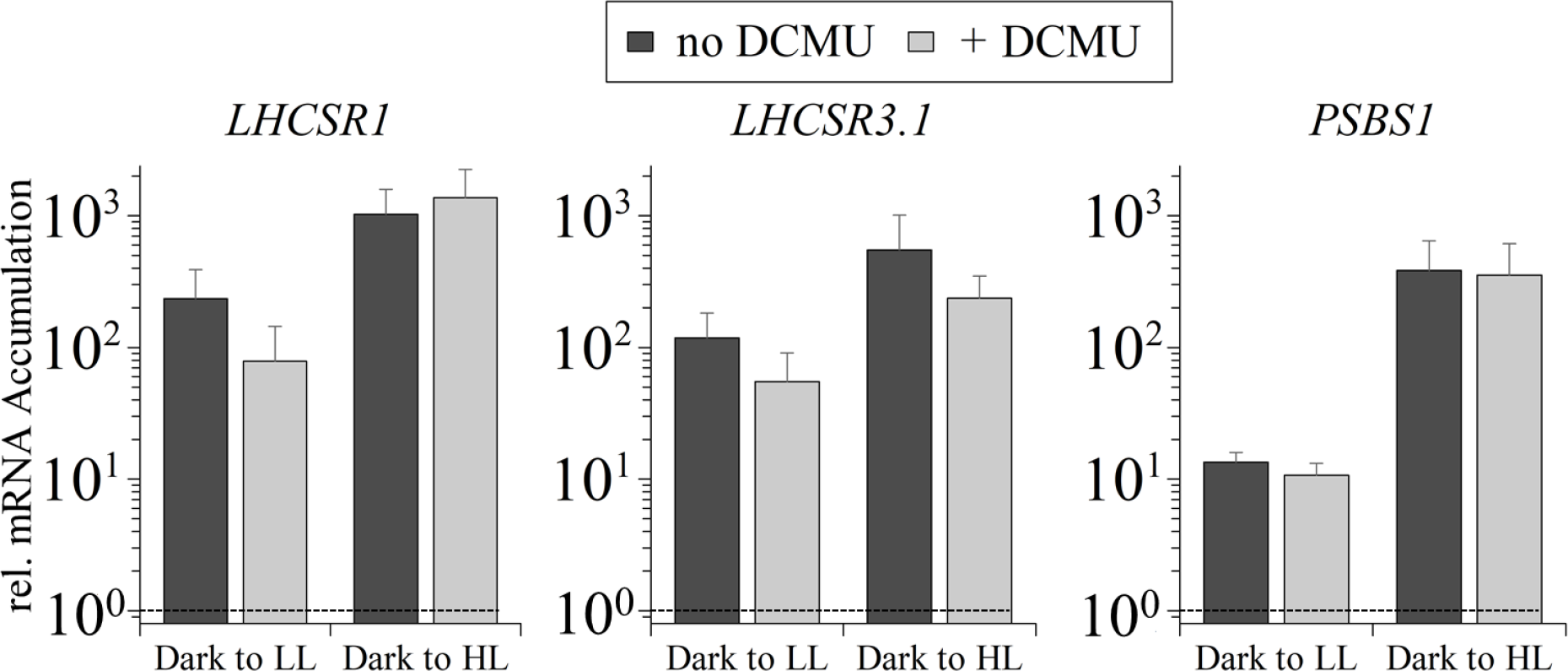
Abundances of *LHCSR1*, *LHCSR3.1* and *PSBS1* transcripts after one hour in LL and HL, in the absence and presence of DCMU. WT CC125, grown as described in the legend of **Figure 1**, was exposed to either white LL (30 µmol photons m^-2^ s^-1^) or HL (480 µmol photons m^-2^ s^-1^) for 1 h, in the absence or presence of 10 µM DCMU, which was added to the cultures immediately before light exposure. Data were normalized to 1 (shown as dotted line in graph), which was set as the initial dark level of the mRNA. There was no significant difference of transcript levels comparing the same samples treated with and without DCMU (although slight differences are seen in this graph). Accumulation of *PSBS1* transcripts between LL and HL was highly significant (p≤0.0001), as well as the *LHCSR1* transcript in cells treated with DCMU in LL and HL (p=0.0153). n=3+SD. Statistical analyses and P-values are listed in **Supplementary Table 2.**

To elucidate the importance of the pre-acclimation conditions on induction of the photoprotective genes and explore differences between our results and those obtained previously (Petroutsos et al., 2016), *LHCSR3.1* transcript levels were quantified in WT cells, either acclimated to the dark or LL (15 µmol photons m^-2^ s^-1^) in High Salt Medium (HSM) overnight and then transferred to HL (300 µmol photons m^-2^ s^-1^) for 1 h either in the presence or absence of DCMU. In the LL acclimated cells, transcript accumulation in HL was completely inhibited by the addition of DCMU (**Supplementary Figure 2**). However, after a pre-incubation in the dark, DCMU-treated cells exposed to HL exhibited an increase in the level of *LHCSR3.1* of 44-fold (**Supplementary Figure 2**), which is in accord with the results presented in **Figure 2**.

These data highlight the impact of the pre-acclimation conditions (those under which cells are maintained prior to the test conditions) on transcript levels. Transcript levels may be much lower in cells kept in dark relative to LL conditions, which would make the fold induction in HL much higher in cells coming from dark relative to LL. In addition, a transition from dark to HL may be more stressful than from LL to HL since, in the latter, the PET system is already active and coupled to CO_2_ fixation (the Calvin-Benson-Bassham cycle, CBBC) while in the former CO_2_ fixation would have to be activated following the dark incubation and the initial exposure to HL might generate more reactive oxygen and nitrogen species (together referred to as RS) that could stimulate activation of the photoprotective genes.

### PHOT1 regulates initial low light responses

After showing that LL is sufficient to cause substantial accumulation of *LHCSR1*, *LHCSR3.1* and *PSBS1* mRNA, we tested whether this LL induction was blue- and/or red-light dependent. *LHCSR1*, *LHCSR3.1* and *PSBS1* transcripts were quantified following exposure of WT cells to low levels of blue, red and white light (see spectra of light sources in **Supplementary Figure 3**). For WT cells, transcripts from the three photoprotective genes increased 2-3 orders of magnitude relative to control cells when exposed to blue light (30 µmol photons m^-2^ s^-1^), as shown in **Figure 3A** and **3B**. A similar level of transcript accumulation was observed in white light (30 µmol photons m^-2^ s^-1^) (**Figure 3A**). Red light exposure (30 µmol photons m^-2^ s^-1^) led to much lower transcript accumulation than in either blue or white light (**Figure 3A**, note log scale). Changes in levels of transcripts from the *LHCSR3.2* and *PSBS2* genes in response to blue and red light were similar to those observed for *LHCSR3.1* and *PSBS1*, respectively (**Supplementary Figure 4A**).

**Figure 3:**
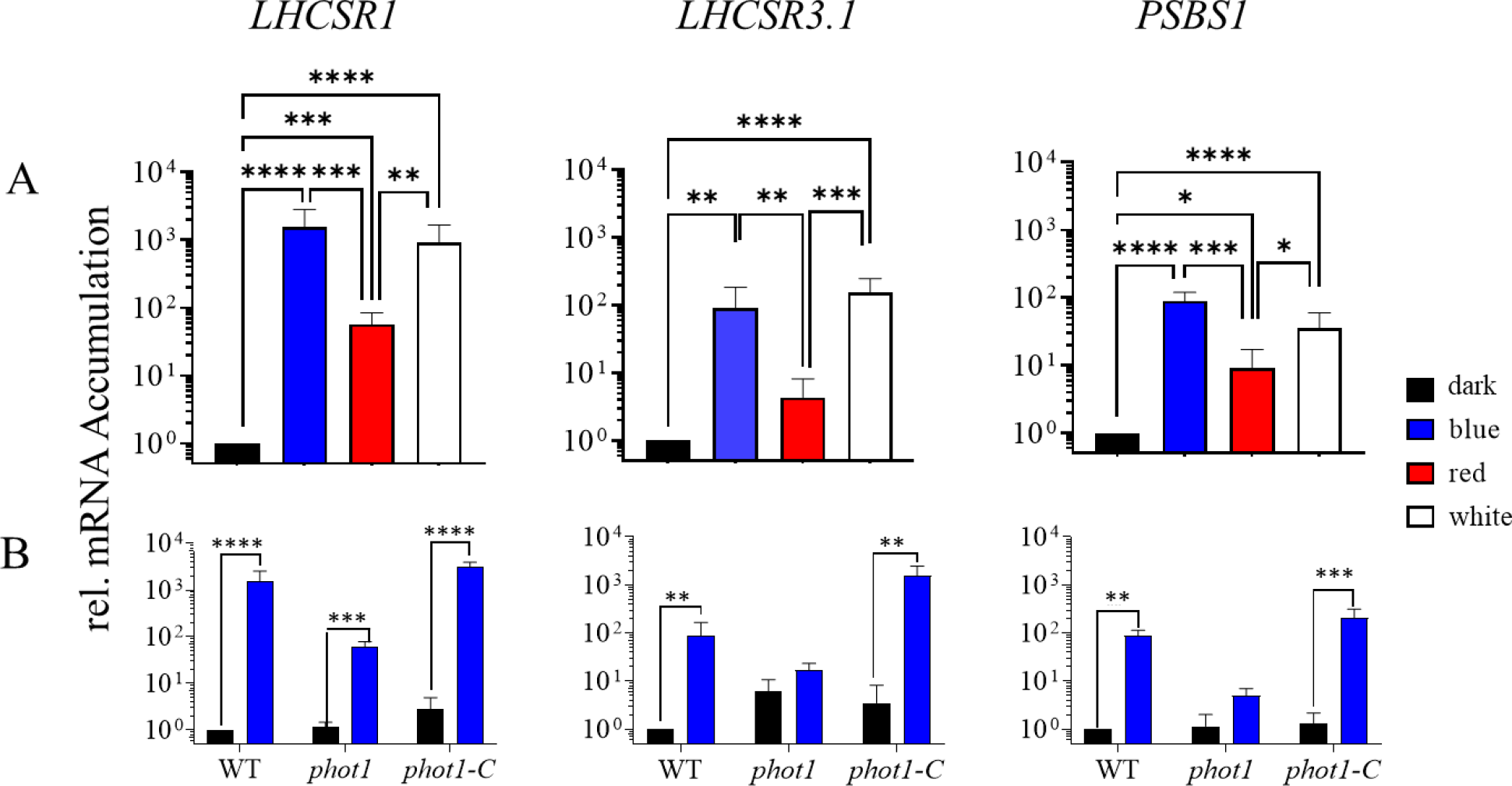
Transcript accumulation of *LHCSR1, LHCSR3.1* and *PSBS1* after exposure to blue, red or white light. WT (CC-125), *phot1* and *phot1-C* cells were grown as described in the legend of **Figure 1. A)** WT was induced with 30 µmol photons m^-2^ s^-1^ blue (blue bar), red (red bar) or white (white bar) light for 1 h. **B**) WT, *phot1* and *phot1-C* cells were induced as in (**A**). All transcript levels were normalized to that of the WT in the dark. n=3-5+SD. Statistical analyses and P-values are listed in **Supplementary Table 3.**

The similar level of transcript accumulation in blue and white light and the much weaker impact of red light suggests that most of the LL increase in expression of these genes is explained by the impact of the blue light photoreceptor PHOT. To substantiate this, blue light dependent transcript accumulation was examined in WT cells, the *phot1* mutant and the PHOT1 rescued strain (*phot1-C*). The *phot1* mutant used in this analysis was generated by CRISPR-Cas9 editing (Greiner et al., 2017) and exhibits a similar phenotype to that of the previously published *phot1* mutant (Petroutsos et al., 2016); there is a loss of the PHOT1 protein, a reduction in the level of the LHCSR3 protein (**Supplementary Figure 5A**) and a marked decrease in the capacity of the cells to perform non-photochemical quenching (NPQ) (**Supplementary Figure 5B**). In the *phot1-C* strain, *PHOT1* is ectopically expressed, reaching a transcript level similar to that of WT cells; this strain has a restored capacity to synthesize high levels of LHCSR3 protein and to perform NPQ (**Supplementary Figure 5)**. As shown in **Figure 3B**, the blue light triggered increases in the transcripts from the photoprotective genes were strongly suppressed in the *phot1* mutant and rescued in the *phot1-C* strain. *LHCSR3.1* and *PSBS1* showed very little mRNA accumulation (not statistically significant) in low blue light-exposed *phot1* cells whereas the *LHCSR1* transcript still exhibited a low, but significant level of accumulation in the mutant. Overall, disruption of the *PHOT1* gene led to a ≥96% reduction in accumulation of all photoprotective gene transcripts following exposure to 30 µmol photons m^-2^ s^-1^ blue light (**Figure 3B**).

These results demonstrate that LL-elicited accumulation of the photoprotective transcripts is strongly dependent on blue light photoperception by PHOT1, with potentially a small impact through an alternative photoreceptor (e.g., cryptochromes) and/or a photoreceptor-independent pathway that is responsive to both red and blue light. The PHOT1-independent light effect on accumulation of these transcripts, especially for *LHCSR1*, could also reflect a small impact of electron transport in modulating accumulation of these transcripts, which is supported by the finding that DCMU impacts their abundances to a small extent (especially in LL), as noted in **Figure 2** and **Supplementary Figure 4B.**

### UV-B radiation elicits UVR8-dependent, PAR independent, accumulation of mRNA from the photoprotective genes

As shown in **Figure 1A**, UV-B light peaks at the same time of the day as PAR, although its increase is delayed relative to PAR and its decrease occurs several hours ahead of the PAR decrease, with very low intensity during the early morning and late afternoon. Furthermore, unlike PAR, UV-B is not diminished much by cloud cover. The role of UV-B radiation on expression of the photoprotective genes was previously investigated, showing that supplementation of very low light (5 µmol photons m^-2^ s^-1^) with UV-B radiation leads to an increase in accumulation of *LHCSR1*, *LHCSR3.1* and *PSBS1* transcripts (Allorent et al., 2016). Similarly, we observed that UV-B light has an augmenting effect on transcript accumulation when cells were exposed to 30 µmol photons m^-2^ s^-1^ white light (**Supplementary Figure 6**). For these studies we used two WT strains (CC-124 and CC-125) and monitored the kinetics of transcript accumulation following exposure of the cells to LL and LL+UV-B radiation over a 1 h period. A gradual accumulation of each of the three transcripts was observed, with a significant difference (10-20-fold) between LL and LL+UV-B after 1 h of irradiation (**Supplementary Figure 6**).

The kinetics of induction of the target genes in CC-124 were slightly different relative to CC-125; the transcripts reached maximal levels a little more rapidly in CC-124. The maximum difference between the levels of these transcripts measured in LL and LL+UV-B appeared to occur between 30 min and 1 h. *PSBS1* transcript accumulation appeared to be more strongly elevated in CC-124 by supplementation with UV-B radiation than that of *LHCSR1* or *LHCSR3.1*, especially when measured shortly after the initiation of UV-B exposure (15 min), reaching a maximal level after approximately 30 min, which agrees with previously published data (Allorent et al., 2016). Overall, supplementation of LL maintained cells with UV-B radiation caused a marked (≥10-fold) increase in levels of mRNA from the three photoprotective genes after 15-60 min of UV-B exposure.

The UV-B radiation used in these experiments (200 µW/cm^2^) corresponds to the maximum intensity observed at noon on a summer day in California (**Figure 1A**). Usually, this UV-B level is accompanied by the highest PAR intensity measured during the day, although at times, much of the PAR can be blocked by cloud cover without strongly impacting UV-B penetrance. To dissect the specific contribution of UV-B light, we examined its effect on gene expression in the presence or absence of high PAR. Additionally, mRNA accumulation was measured in both WT cells and the *uvr8* mutant, which is null for the UV-B photoreceptor. As shown in **Figure 4A**, UV-B irradiation of WT cells in the absence of PAR elicited a surprisingly high increase in accumulation of *LHCSR1*, *LHCSR3.1* and *PSBS1* transcripts. The extent of this increase for the *LHCSR1* and *PSBS1* transcripts was essentially identical to that observed when the cells were exposed to HL (480 µmol photons m^-2^ s^-1^), with no additional increase in cells simultaneously exposed to HL plus UV-B radiation. However, the highest level of *LHCSR3*.*1* transcript accumulation occurred in cells exposed to both HL and UV-B radiation (increase by an additional ∼5 fold with UV-B irradiation). This observation may reflect an inability of the levels of either UV-B or HL radiation alone to fully saturate the induction of *LHCSR3.1*; the inability to saturate the *LHCSR3.1* transcript accumulation at 480 µmol photons m^-2^ s^-1^ was also observed in **Figure 1** where the level of the *LHCSR3.1* mRNA under the highest irradiation, 960 µmol photons m^-2^ s^-1^, was elevated by 2-3-fold relative to the level at 480 µmol photons m^-2^ s^-1^. Therefore, we performed the same experiment as in **Figure 4**, but the PAR light level used was 960 µmol photons m^-2^ s^-1^ (very high light, VHL) (**Supplementary Figure 7A**). Similar to the observations presented in **Figure 1**, *LHCSR3.1* transcript accumulation in cells exposed to VHL was about twice as high as that of cells exposed to HL, with a similar level attained when the cells were only exposed to UV-B radiation. However, transcript accumulation for VHL and HL both supplemented with UV-B followed the exact same trend (compare **Supplementary Figure 7A** with **Figure 4**); the level of the *LHCSR3.1* transcript was identical in UV-B and in VHL, while combining the two light sources led to higher *LHCSR3.1* transcript accumulation (∼5-fold). These results suggest that while PAR and UV-B light can reach similar levels and compensate for each other with respect to *LHCSR3.1* mRNA accumulation, only simultaneous exposure to both types of radiation promote maximum transcript accumulation under the PAR levels tested in this work.

**Figure 4:**
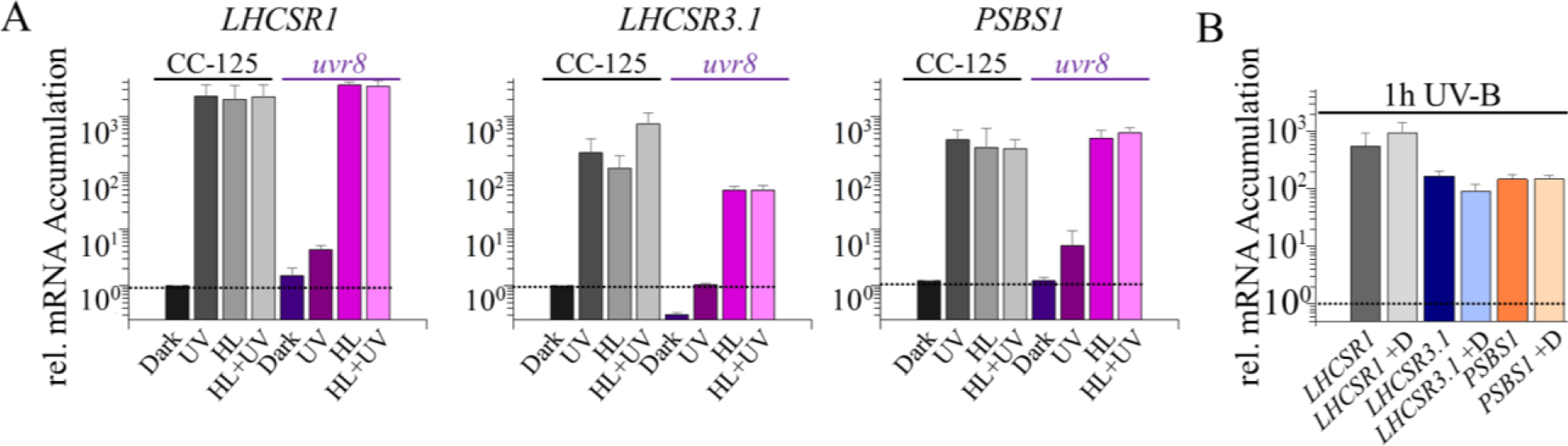
Impact of UV-B radiation on expression of photoprotective genes. **A**) Changes in levels of *LHCSR* and *PSBS* transcripts after 1 h of UV-B (UV) radiation, HL, or HL+UV-B radiation. WT CC-125 (grey-black bars) and *uvr8* cells (coloured bars) were grown as described in the legend of **Figure 1**. Cultures were then divided and either exposed to UV-B irradiation (200 µW/cm^2^), exposed to HL (480 µmol photons m^-2^ s^-1^) or to HL in the presence of UV-B radiation. **B**) Cultures were grown as in **A** and all samples were exposed to UV-B radiation in either the absence or presence of 10 µM DCMU (+D). The dotted line across the bar graph indicates the level of transcript in the dark before illumination. All transcript levels in both **A** and **B** were normalized to the transcript level of the WT in the dark. n=3-7+SD. Statistical analyses and P-values are listed in **Supplementary Table 4.**

We next tested the role of the UVR8 photoreceptor in controlling accumulation of transcripts from the photoprotective genes. The *uvr8* mutant had markedly reduced levels of transcripts from all three of the photoprotective genes relative to WT cells following UV-B exposure (**Figure 4A**), indicating that UVR8 is integral to the regulation of these transcripts. For *LHCSR1* and *PSBS1*, the transcript levels attained in HL and HL+UV-B radiation were not affected by the loss of the UVR8 photoreceptor. However, the mutant exhibited slightly lower levels of *LHCSR3.1* mRNA in HL and HL+UV-B radiation; this reduction was 3-4-fold relative to WT cells, although the overall fold induction was comparable in both strains (the dark levels of the *LHCSR3.1* transcript were lower in the *uvr8* mutant); the differences are not statistically significant. Similar results were obtained for the impact of HL and HL+UV-B radiation on the patterns of transcript accumulation for *LHCSR3.2* and *PSBS2* relative to those of *LHCSR3*.*1* and *PSBS1*, respectively (**Supplementary Figure 7B**). Additionally, as shown in **Figure 4B**, the addition of DCMU had no impact on UV-B induced expression of these genes, demonstrating that all transcript accumulation during exposure to UV-B is independent of linear photosynthetic electron transport.

This data also suggests that there is a small UV-B dependent, UVR8-independent accumulation of the *LHCSR1* and *PSBS* transcripts (**Figure 4A**). Since the *uvr8* mutant lacks the UV-B photoreceptor and the transcript levels are not impacted by DCMU, we suggest that this effect could be triggered by reactive species (RS) such as ROS, which can be generated as a consequence of UV-B radiation-mediated damage (de Jager et al., 2017; Rastogi et al., 2014; Yokawa et al., 2016), or be a consequence of stimulation of the PHOT1 photoreceptor which has very low absorption in the UV-B region of the spectrum.

### Light intensity and CO_2_ levels independently impact accumulation of transcripts from the photoprotective genes

CIA5 is a regulatory element that controls acclimation of Chlamydomonas to low CO_2_ conditions [e.g., induction of carbon concentrating mechanism (CCM)] (Fang et al., 2012; Xiang et al., 2001). Although *LHCSR3* gene expression has been associated with HL, it was also shown to be highly dependent on the level of CO_2_; transcript levels decreased when the CO_2_ concentration of the culture was elevated (Wang et al., 2016). Furthermore, previous work reported that expression of *LHCSR3* is impacted in the *cia5* mutant (Fang et al., 2012).

To study the role of CIA5 in the regulation of the photoprotective genes and determine whether the light- and CO_2_-CIA5-dependent transcriptional regulation of these genes are linked, WT, the *cia5* null mutant, and a *cia5* rescued strain (*cia5* mutant with wild-type *CIA5* gene ectopically expressed; *cia5-C*) were exposed to LL, moderate light (120 µmol photons m^-2^ s^-1^, ML), and VHL (1000 µmol photons m^-2^ s^-1^) at both ambient and high CO_2_ levels and changes in *LHCSR1*, *LHCSR3.1* and *PSBS1* transcript levels were analyzed. As shown previously, the absence of CIA5 negatively impacted accumulation of the *LHCSR3*.*1* transcript (Fang et al., 2012; Wang et al., 2016), however, our results also show that VHL intensities can partially compensate for the lack of CIA5 as differences in transcript accumulation were smaller between WT and *cia5* strains when the light intensity was increased (**Figure 5A**). The *cia5* mutant exhibited a significant increase in *LHCSR3.1* transcript accumulation at ML and VHL; in VHL the mRNA accumulation was ∼15 fold higher than in LL and >100 fold higher than in the dark. This mRNA accumulation in the mutant supports the idea that light can regulate *LHCSR3.1* expression in a CIA5-independent way. Furthermore, elevated CO_2_ levels strongly suppressed transcript accumulation leading to similar *LHCSR3.1* mRNA levels in both the WT and *cia5* strain, since in high CO_2_ conditions, there is no requirement/role for CIA5 regulation. Nevertheless, the *LHCSR3.1* gene was still induced under high CO_2_ concentrations at the higher light intensities, also pointing to the participation of a CIA5-independent pathway in *LHCSR3.1* transcriptional regulation. Moreover, *LHCSR3.1* transcript accumulation was also analyzed in WT, *phot1* and *cia5* in blue LL (**Supplementary Figure 4D**), which demonstrated that the mRNA levels in WT and *cia5* showed the same trend in blue as in white LL (**Figure 5**). Interestingly, the PHOT1 dependent increase in *LHCSR3.1* transcript abundance was strongly suppressed by high CO_2_ (**Supplementary Figure 4D**), which raises the possibility that the PHOT1-dependent regulation might be an indirect effect caused by a reduction in the CO_2_ levels. In contrast to the results obtained for *LHCSR3.1*, CIA5 barely affected *LHCSR1* expression in cells exposed to LL, ML or VHL in the presence or absence of 5% CO_2_ (**Figure 5**), while *PSBS1* transcript accumulation in the absence of CO_2_ supplementation in LL, ML and VHL was similar in WT, *cia5* and the *cia5-C* rescued strain (see below for further discussion).

**Figure 5:**
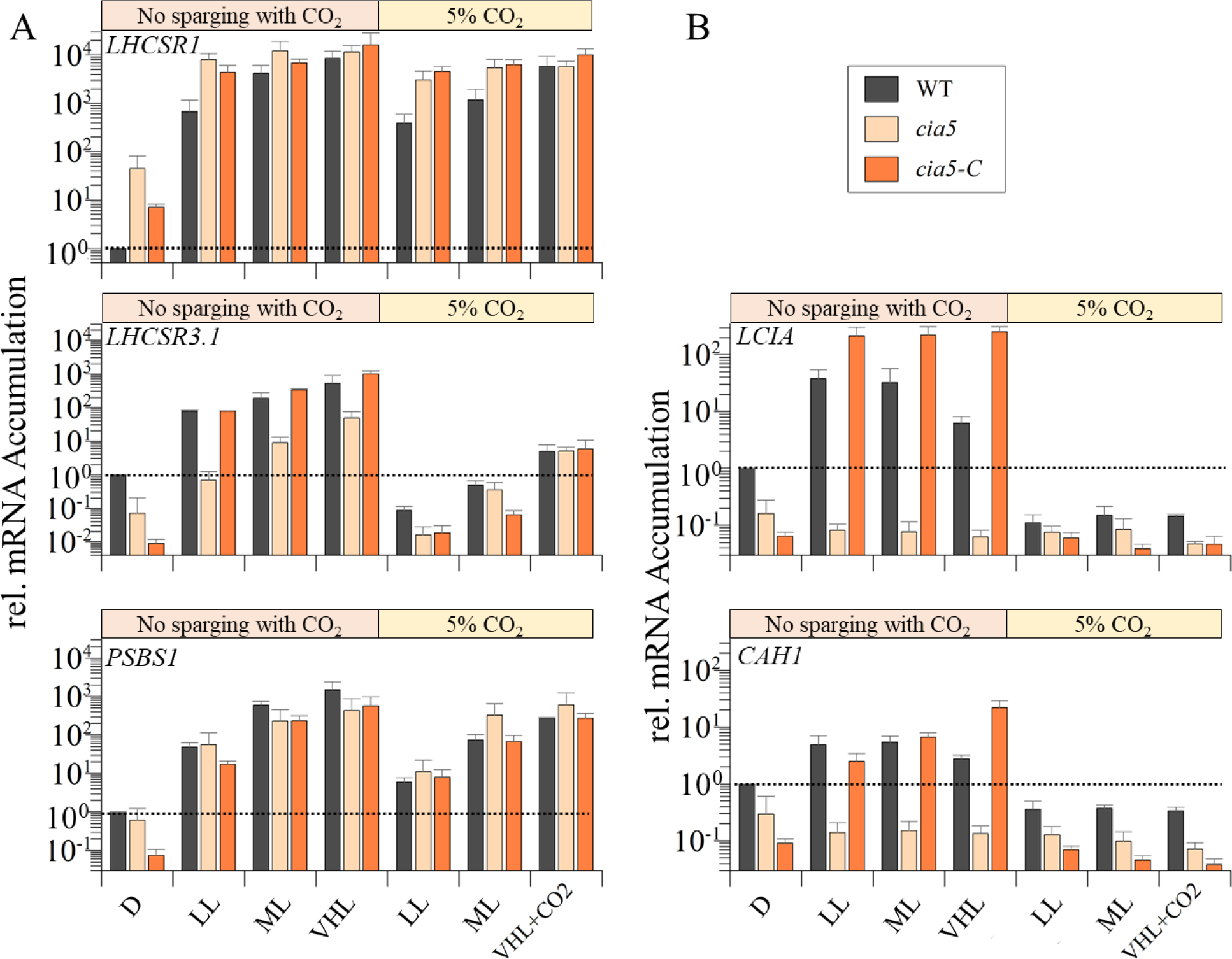
Impact of different light intensities and CO_2_ levels on expression of photoprotective genes. CC-125, *cia5* mutant cells and the rescued *cia5-C* strains were grown and dark adapted as described in the legend of **Figure 1** and then transferred to HSM or HSM +5 % CO_2_ for 1h at either LL (30), ML (120) or VHL (1000 µmol photons m^-2^ s^-1^). Transcript levels were normalized to the level in WT cells prior to induction (D, dotted line). n=3+SD. Statistical analyses and P-values are listed in **Supplementary Table 5.**

We performed the same analyses as described above with two CCM genes that were previously shown to be under CIA5 control (Tirumani et al., 2014). These genes (*CAH1* and *LCIA*) were upregulated at ambient CO_2_ levels, in the presence of light, and in a CIA5-dependent way in WT cells. Contrary to the results for *LHCSR3.1*, *CAH1* and *LCIA* transcript accumulation was similar under all light conditions used in this experiment (LL, ML and VHL) independent of the CO_2_ regime. Their induction was suppressed in WT cells sparged with 5% CO_2_ (under all light conditions), not observed in the *cia5* mutant at any light intensity, and the phenotype was rescued in the complemented strain, which often exhibited even higher transcript levels than the WT strain (possibly due to overexpression of the ectopic *CIA5* gene) (**Figure 5B**). These results suggest that CIA5 is absolutely required for expression of *CAH1* and *LCIA* and the light effect is probably caused by a reduction in CO_2_ levels. Overall, our results confirm that CIA5 is essential for expression/induction of CCM genes (*CAH1* and *LCIA*) under low CO_2_ conditions while it appears to function as an enhancer for *LHCSR3.1*, which can still be upregulated by CIA5-independent light-dependent signals.

### Integration of CO_2_ and UV-B light signals in the transcriptional regulation of the photoprotective genes

We also tested whether UV-B-elicited responses in transcript abundances for the photoprotective genes were linked to CO_2_ concentrations and CIA5 regulation. We exposed WT, *uvr8* and *cia5* strains to UV-B light with or without 5% CO_2_, and measured transcript accumulation for the three photoprotective and the two CCM genes previously studied. Interestingly, the UV-B dependent 400-fold accumulation in *LHCSR3.1* transcript observed in WT cells was completely abolished by sparging the culture with 5% CO_2_ (**Figure 6A**). However, the *cia5* mutant still exhibited an increase in *LHCSR3.1* transcript accumulation following UV-B radiation by almost two orders of magnitude (comparable fold change to WT). Nevertheless, the overall transcript levels were highly depressed in the dark and after UV-B light exposure in the mutant compared to the WT strain. These results suggest that *LHCSR3.1* upregulation mediated by UV-B is CIA5-independent, but also that CIA5 acts either directly or indirectly to enhance the overall expression of this gene in the dark and during exposure to UV-B radiation (**Figure 6A**). The lower mRNA levels in the *cia5* mutant strain in the dark indicate that *LHCSR3.1* expression was already induced in the dark in a CIA5-dependent way in the WT strain. In our experiments, dark-acclimated cells were transferred from TAP to acetate free medium (HSM) 2 h before the UV-B exposure. We measured *LHCSR3.1* transcript levels before and after transferring WT and *cia5* cells to HSM and we could see that this transfer led to *LHCSR3.1* mRNA accumulation (**Figure 6B**). This result can be explained based on the recent findings of Ruiz-Sola and collaborators who demonstrated that changes in CO_2_ availability can activate *LHCSR3* gene expression even in the absence of light in a CIA5-dependent manner (Ruiz-Sola et al., 2021). In our case the transfer of the cells from TAP to HSM would drop CO_2_ levels in the culture [CO_2_ that accumulated in TAP medium (associated with metabolism of acetate) in the dark would decline] and therefore activate *LHCSR3*. Under high CO_2_ conditions, CIA5 would not be active and, as observed in white light, *LHCSR3.1* exhibited similar transcript levels in both WT and *cia5* strains.

**Figure 6:**
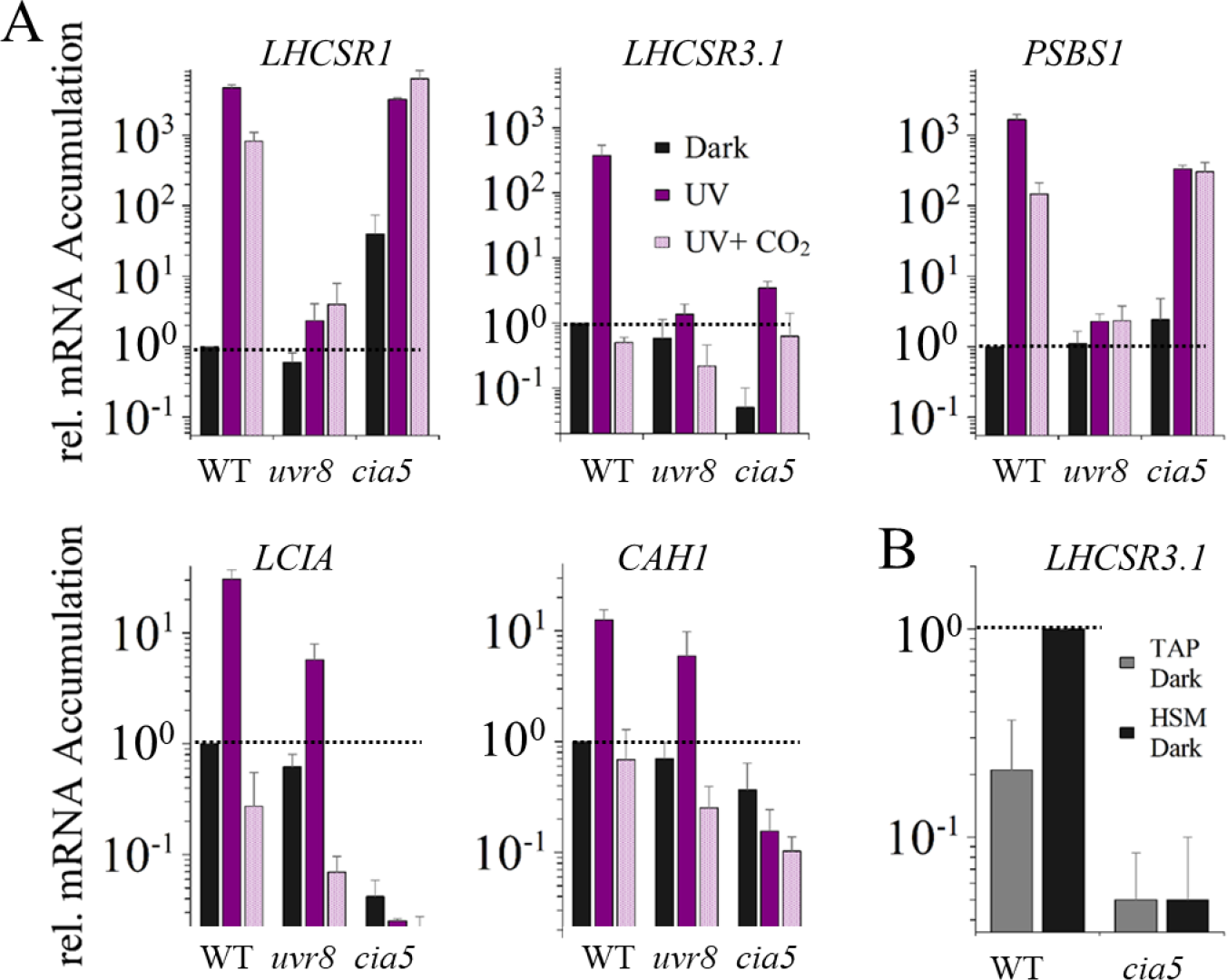
Impact of UV-B radiation, CO_2_ and CIA5 on expression of CCM genes and photoprotective genes. Changes in levels of transcripts from **A)** the CCM genes *LCIA* and *CAH1* and the photoprotective genes *LHCSR1*, *LHCSR3*.1 and *PSBS* after 1 h of UV-B radiation (200 µW/cm^2^) in the absence (UV, dark purple bars) or presence (UV+ CO_2_, faded purple bars) of 5% CO_2_ in the WT, *uvr8* and *cia5* strains; and **B)** from *LHCSR3.1* in the dark after 24 h in TAP (grey) and after 2 additional hours following the change of the culture from TAP to HSM (black) in the WT and *cia5* mutant. n=3+SD. Transcript levels were normalized to the initial level of WT in the dark. Statistical analyses and P-values are listed in **Supplementary Table 6.**

Interestingly, the CCM-related genes exhibited a different regulation compared to *LHCSR3.1*; *LCIA* and *CAH1* mRNA accumulation in cells exposed to UV-B was similar to that observed under white light (**Figure 5**) and strongly repressed by CO_2_ but, contrary to *LHCSR3.1*, this induction was not UVR8-dependent (it was between 10- and 12-fold for both WT and *uvr8*). Furthermore, unlike *LHCSR3.1*, *LCIA* and *CAH1* transcript accumulation was completely abolished in the *cia5* mutant under all conditions. This indicates that these genes might be exclusively regulated by the CO_2_/CIA5 signaling pathway and suggests that the light-mediated responses (PAR and UV-B) may be indirect, altering the level of CO_2_ and the efficacy of the CO_2_/CIA5 signaling pathway. Understanding whether UV-B light can alter the intracellular CO_2_ levels or whether the CCM genes might respond to changes in RS mediated by UV-B radiation will require further investigation.

While *LHCSR1* and *PSBS1* transcripts strongly accumulated upon exposure to UV-B radiation (as shown in **Figure 4**), when the cultures were sparged with 5% CO_2_ concomitant with the UV-B exposure, there was only a 5-fold decrease of the *LHCSR1* transcript and a 12-fold decrease of the *PSBS1* transcript, indicating that CO_2_ does not have a strong impact on expression of these genes in dark-to-light transitions. Furthermore, *LHCSR1* and *PSBS1* transcript levels still increased in *cia5* cells induced by UV-B light to a level similar to that observed in WT cells and this induction was not suppressed by sparging the cultures with high CO_2_. Hence, unlike for the *LHCSR3.1* transcript, CIA5 has little impact on expression of the *LHCSR1* and *PSBS1* genes during UV-B dependent induction.

## DISCUSSION

The fastest NPQ mechanisms induced under HL are thermal dissipation (qE) and state transition (qT), which become active in the range of seconds (qE) to minutes (qT) (Nilkens et al., 2010; Takahashi et al., 2013). While the transcript levels of the qE related genes are extremely low and the proteins are absent in dark-acclimated cells, their induction must anticipate HL exposure to minimize cellular damage. In this work, we analyzed how photosynthetic cells sense and integrate environmental cues to modulate expression of the photoprotective genes *LHCSR1*, *LHCSR3.1* and *PSBS1*, and the adaptation mechanisms that they have incorporated for coping with light stress prior to accumulation of the encoded proteins. We show that illumination with even very LL was sufficient to cause substantial accumulation of transcripts from the photoprotective genes in dark pre-acclimated cells (**Figure 1B**). This LL induction was mainly mediated by blue light and required PHOT1 as cells deficient in PHOT1 exhibited a marked reduction in accumulation of *LHCSR1, LHCSR3.1* and *PSBS1* transcripts (**Figure 7**), as it has been shown before via RT-PCR (Aihara et al., 2019). Thus, at dawn, with the first low levels of sunlight, Chlamydomonas would induce the photoprotective genes. Sun light is blue light-enriched in aquatic environments, as shorter wavelengths in a water column have a higher penetrating capacity (Kirk, 1994). Furthermore, the proportion of the blue light in the visible spectrum reaching the Earth’s surface increases from dawn to mid-day and decreases towards sunset. The low irradiance of blue light required to induce photoprotective genes would make this signaling system effective at priming NPQ in both terrestrial and aquatic organisms over the course of the day, in habitats that are mostly light exposed or shaded.

**Figure 7:**
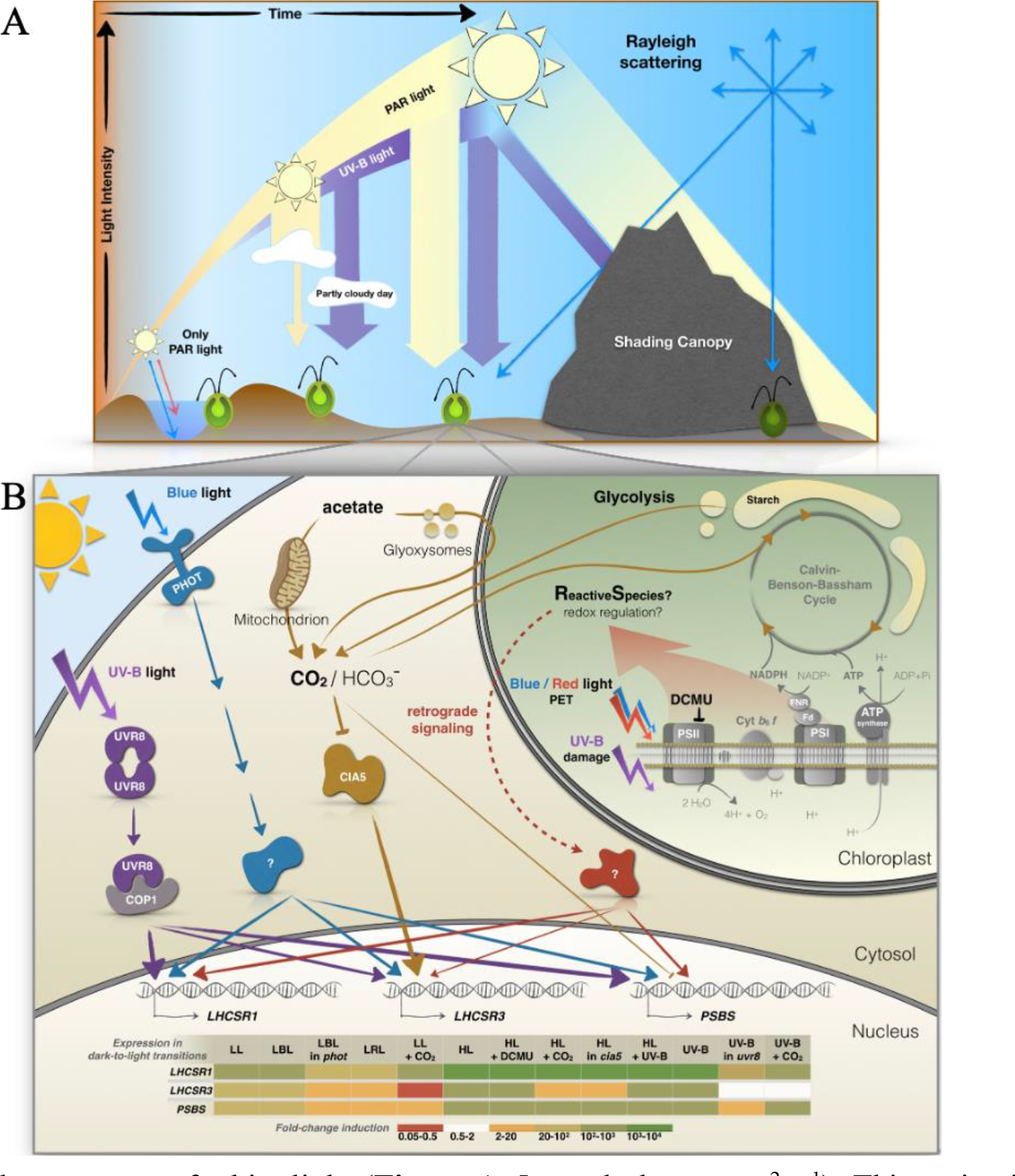
A) Schematic summary of changing light quality and quantity throughout the day. Light intensities increase from dawn to noon. Blue light reaches deeper levels of the water column, while red light is absorbed near the water surface. Additionally, PAR can be strongly reduced by cloud cover, while UV-B radiation might even increase on partly cloudy days. PAR and UV-B radiation is strongest during noon. While direct sunlight is shielded/reduced by canopy shading, blue light (and also UV light in the case of a plant canopy) reach shaded areas more effectively than other wavelength of PAR through Rayleigh scattering, which increases as the wavelength of light decreases. **B) Signals that regulate energy dissipation in Chlamydomonas.** Transcription of *LHCSR1*, *LHCSR3* and *PSBS* is strongly initiated with exposure to a very low amount of white light (**Figure 1**, 5 µmol photons m^-2^ s^-1^). This activation is strongest for *LHCSR3* but also is also apparent for *LHCSR1* and *PSBS* and is dependent on the Chlamydomonas blue light dependent photoreceptor PHOT1. All three transcripts are also partially regulated by PET downstream of PSII and the generation of retrograde signals by HL (red). UV-B exposure further regulates transcript induction on several levels. UV-B radiation directly facilitates monomerization of the UVR8 homodimer, which then binds to COP1 and allows the participation of other factors (not included in the figure) in the transcriptional regulation of the photoprotective genes (purple). UV-B exposure may also lead to the generation of RS in the chloroplast which further triggers signaling events (red). Additionally, *LHCSR3* is strongly controlled by CO_2_ levels and the regulatory factor CIA5, which may function as an enhancer while *PSBS* may be affected by CO_2_ to a minor extent (orange). Additional discussion of the role of CO_2_ in regulating *LHCSR3* is presented in Ruiz et al (Ruiz-Sola et al., 2021). The heatmap table summarizes the transcript fold change for each gene in the transition from dark to the indicated conditions.

However, blue light is only one signal of a complex process of control associated with quenching. In addition to PHOT1-mediated responses, our data suggest that other signals impact photoprotective gene induction in LL, but that the genes have different sensitivities to some of these signals. For example, *LHCSR1* transcript levels exhibited a low but significant increase in the *phot1* mutant when exposed to blue light (**Figure 3B**), reaching levels almost identical to those observed in red light-induced WT cells (**Figure 3A**). Similar behavior, although much less pronounced, was observed for *LHCSR3.1* and *PSBS1*. This low level PHOT1-independent induction could be mediated either by signals generated as a consequence of electron flow in the chloroplast and/or by other photoreceptors such as cryptochromes (CRYs), which can be activated by both blue and red light (Beel et al., 2012). CRYs accumulates during the dark period of the diel cycle, and elicits activation of cellular processes after illumination (e.g., gametes formation, circadian regulation) (Müller et al., 2017). Moreover, they are required for LHCSX protein accumulation in diatoms (Juhas et al., 2014) linking CRYs and photoprotection in microalgae. Integration of CRY- and PHOT1-dependent pathways might enhance expression of the photoprotective genes. Furthermore, CRYs are also required for phytochrome signaling functions associated with the circadian clock in plants (Devlin & Kay, 2000). Although currently there are no identified phytochromes in Chlamydomonas, there are red light stimulated processes in this alga and it has the capacity to synthesize bilins (linear tetrapyrroles that often absorb red light), which may function as signaling molecules and control the biosynthesis of chlorophyll (Duanmu et al., 2013; Zhang et al., 2021). The Chlamydomonas genome also encodes proteins with potential bilin binding domains, like those of phytochromes. The role of CRYs and bilin-based photoreceptors in induction of photoprotective mechanisms in Chlamydomonas requires further investigations.

While photoreceptors may be responsible for increased expression of the photoprotective genes after cells in sustained darkness are exposed to light, these genes would already be active in cells maintained in LL; their transcripts would be higher relative to cells maintained in darkness. The increased basal transcript levels of the photoprotective genes in LL pre-acclimated cells, due to PHOT1, would reduce the apparent impact of exposure to higher light conditions and increase the apparent impact of DCMU if a portion of the light dependent transcript induction was the result of photosynthetic electron flow.

Signals generated by PET that impact gene expression would include the accumulation of RS and redox signaling, especially as the light intensity increases (**Figure 7**, retrograde signaling). Blocking PSII-dependent PET with DCMU led to a slight reduction in *LHCSR3.1* transcript accumulation when the dark pre-acclimated cells were transferred to both LL and HL. Similarly, *LHCSR1* was also mildly repressed by DCMU in LL. Additionally, a small DCMU-dependent repression was observed in low blue light (**Supplementary Figure 4B**), confirming the occurrence of a small (but consistent) DCMU effect, and demonstrating that PSII-dependent electron transport can impact the activities of the *LHCSR* promoters. Unlike *LHCSR1* and *LHCSR3.1*, *PSBS1* mRNA accumulation was not significantly affected by DCMU at any of the light intensities or under either of the pre-acclimation conditions used.

Regarding how PET affects expression of the photoprotective genes, previous studies have shown that the redox state of the PQ pool is not relevant for this regulation. Two photosynthesis inhibitors, DCMU, which promotes oxidation of the PQ pool, and DBMIB, which leads to overreduction of the PQ pool, inhibit LHCSR3 protein accumulation (Petroutsos, Busch, Janßen, et al., 2011). Singlet oxygen production, which is mainly synthesized in the PSII antenna, should be enhanced in the presence of both inhibitors, suggesting that this RS most likely does not cause induction of the *LHCSR3* or *LHCSR1* genes. Furthermore, DBMIB blocks electron transport at the Q_o_ site of the Cyt *b_6_f* complex. Therefore, the signal required for this electron transport-dependent induction of the photoprotective genes is likely generated downstream of the Q_o_ site. Moreover, dark pre-acclimated cells are in state 2 (mobile PSII antenna associated with PSI) due to a reduction in intracellular oxygen levels associated with our experimental conditions (pre-acclimation in the dark without vigorous shaking) (Delepelaire & Wollman, 1985; Finazzi et al., 1999) and the cells would only gradually transition to state 1 in the light, because full activation of the CBBC can take 2-3 minutes (Makowka et al., 2020). Therefore, upon illumination, PSI can absorb more photons because of the larger size of its antenna, which might initially favor RS production because of the initial bottleneck in the CBBC. However, this bottleneck would be ameliorated by the action of the FLV proteins (Chaux et al., 2017) and a diminished supply of electrons from PSII (because of the reduced antennae size). Furthermore, blocking electron flow would prevent the fixation of CO_2_ and lead to an elevation of intracellular CO_2_ levels. Therefore, because of changes in both RS production and accumulation of intracellular CO_2_, it is difficult to define the precise mechanism responsible for reduced expression of *LHCSR3*.*1* in DCMU treated cells.

Under LL conditions, RS generation would not be high enough to elicit the highest levels of *LHCSR1* or *LHCSR3.1* transcripts, even under ambient/low CO_2_ conditions. And while diminished input of electrons to PSI with the addition of DCMU would result in elevated intracellular CO_2_ levels which could negatively impact mRNA accumulation for LHCSR3 (not so much for LHCSR1) (**Figure 2**), the lack of photosynthetic O_2_ evolution would result in lower intracellular O_2_ levels (hypoxia), delayed state 2-to-state 1 transition (Forti & Caldiroli, 2005), elevated cyclic electron flow, the breakdown of starch to generate energy and potentially a tendency to produce RS. In HL, hypoxia and fermentation metabolism may become more prominent, the breakdown of starch and generation of NADH would increase, as would the generation of excited pigment molecules and RS, which could contribute to accumulation of the *LHCSR1* and *LHCSR3* transcripts. Our data suggests that transcription of *LHCSR1* may be more sensitive to RS than *LHCSR3*, which is supported by previous findings (Roach et al., 2020); lower sensitivity of *LHCSR3* expression to chloroplast-generated signals is also supported by the finding that of the three photoprotective genes, this gene required exposure to the highest light intensity (960 µmol photons m^-2^ s^-1^) for maximizing mRNA accumulation (**Figure 1B**).

The discussion above offers explanations for the different induction levels of *LHCSR3* observed in light and dark pre-acclimated cells treated with DCMU (**Supplemental Figure 3**). In contrast to the dark pre-acclimated cells, in cells pre-acclimated in LL the photosynthetic apparatus would be in state 1, PSI would absorb less light energy and photosynthesis and CO_2_ fixation would be coupled from the onset of the experiment. Therefore, RS production from PSI would be lower than in dark pre-acclimated cells with the onset of light, DCMU might further reduce RS production while at the same time promoting a rapid increase in the intracellular CO_2_ levels that would reduce CIA5-dependent accumulation of *LHCSR3* mRNA (as shown in **Figure 5**) without affecting *LHCSR1* expression.

While PHOT1 and potentially other photoreceptors may have an essential role in inducing NPQ-related genes with the very first light (low intensity light) in the morning (blue pathway in **Figure 7**), the chloroplast-generated signals (red pathway in **Figure 7**) would modulate this expression according to light intensity. The higher the intensity, the higher the rate of CO_2_ uptake (until saturation) and the lower the level of internal CO_2_, while more signal is also generated from PET (**Figure 1B**); the highest level of transcript accumulation would reflect both a diminished CO_2_ concentration and elevated production of photosynthetically generated RS.

Several hours after dawn, the solar spectrum is progressively enriched in UV-B (**Figure 1A**). It was previously shown that the three photoprotective genes discussed here are induced in LL+UV-B light (Allorent et al., 2016). However, we have now shown that LL can already significantly induce expression of these genes. Therefore, we studied the capacity of UV-B radiation to induce these genes in the absence of PAR and in the presence of HL. Our results indicate that the levels of the *LHCSR1*, *LHCSR3*, and *PSBS* transcripts in cells solely exposed to UV-B light were similar to those of cells incubated in HL. Moreover, supplementation of HL with UV-B light did not elicit a significant increase in the levels of these transcripts, except for *LHCSR3* which exhibited some increased induction upon exposure to HL+UV-B radiation (**Figure 4**).

The saturating or near saturating response mediated exclusively by UV-B light was strongly diminished in the *uvr8* mutant, although residual low-level induction was still observed for the three genes (**Figure 4**). The increases in the levels of transcripts for *LHCSR1* and *PSBS* in HL in the *uvr8* mutant were essentially the same as those observed in UV-B and HL+UV-B in the WT strain, indicating that PAR and UV-B light act independently and can compensate for each other, achieving the same final output. Furthermore, UV-B light can enhance RS production in both animal and plant cells (de Jager et al., 2017; Gniadecki et al., 2000; Huarancca Reyes et al., 2018; Rastogi et al., 2014; Yokawa et al., 2016), which could drive this residual induction.

The UV-B-dependent, PAR-independent pathway for activation of the photoprotective genes may not represent an advantage under a clear sky, where cells would experience HL prior to being exposed to high levels of UV-B radiation (**Figure 1A**). However, UV-B radiation elicited induction might have a role in maintaining maximal promoter activity for the photoprotective genes during long exposure to HL since the levels of these transcripts peak within the first hour of exposure to HL with a significant reduction during a longer period of HL exposure (Tibiletti et al., 2016). Additionally, UV-B-dependent induction could confer an advantage to the cells when they are under a canopy, which reduces the level of PAR relative to UV-B radiation (Hermanowicz et al., 2019). The perception of UV-B radiation would also help organisms sense the time of day and when the light intensity is likely to be at its highest, even though the organism may not be experiencing excess PAR. This situation is common during a partly cloudy day or in environments in which plants are exposed to intermittent sun-shade periods; cloud cover may dramatically and almost instantaneously reduce the intensity of PAR while having only a minor effect on the intensity of the UV-B radiation (**Figure 1A**). In fact, it has been reported that under partly cloudy skies, UV-B radiation may increase in intensity by ∼25% relative to clear skies (Mims & Frederick, 1994). Thus, perception of UV-B light would allow cells to maintain a primed system for triggering NPQ even when there are variations in the intensity of PAR. It has been shown already that pre-acclimation in LL+UV-B radiation improves survival when the cultures are suddenly exposed to HL (1000 µmol photons m^-2^ s^-1^), and this protection seems to be mainly mediated by LHCSR1 and, to a lesser extent, LHCSR3 (Allorent et al., 2016).

In addition to light intensity and quality, carbon availability also regulates NPQ. It is known that the *LHCSR3* transcript accumulates when inorganic carbon levels are low, and that this induction is mediated by CIA5, the main regulatory factor involved in controlling the genes associated with the CCM (Fang et al., 2012; Polukhina et al., 2016). PSBS protein levels are also elevated more in minimal medium (air levels of CO_2_) than in TAP medium (17 mM acetate) (Correa-Galvis et al., 2016); internal CO_2_ levels would increase in the presence of acetate (Ruiz Sola et al., 2021). In contrast, *LHCSR1* transcript levels accumulated in the presence of high CO_2_, especially upon exposure to HL (Yamano et al., 2008). Our results clearly confirm that *LHCSR3* is dramatically repressed in the presence of high CO_2_ and that the induction under limiting inorganic carbon conditions is strongly regulated by CIA5 (**Figure 5**). However, in this work, we also demonstrate that light impacts *LHCSR3* expression independently of CIA5; the *cia5* mutant responded to different light intensities during exposure to both low and high CO_2_ (**Figure 5**). The higher levels of the *LHCSR3* transcript observed in *cia5* cells incubated under low relative to high CO_2_ could result from the generation of higher levels of RS production in the former as a consequence of CO_2_ depletion and increased use of O_2_ as a terminal electron acceptor. Overall, these results suggest that different signals may converge for controlling the activity of the *LHCSR3* promoter and exert their positive effects independently (**Figure 7**).

We propose that CIA5 acts as an enhancer, a positive regulatory element necessary to potentiate transcriptional regulation in conjunction with other regulatory elements (**Figure 7**). Indeed, we observed that *LHCSR3* was still induced by blue light (**Supplementary Figure 5**) and UV-B radiation (**Figure 6**) in the *cia5* mutant, although both the basal and induced levels of expression were lower than in WT cells (but similar fold-change). The lower *LHCSR3* mRNA levels present in dark-acclimated *cia5* mutant cells are in line with the work by Ruiz-Sola et al. (2021) that demonstrates that CIA5-dependent *LHCSR3* induction also occurred in total darkness when the availability of inorganic carbon becomes very low (**Figure 6B**).

Interestingly, the light signal that regulates *LHCSR3* in the *cia5* mutant did not impact the CCM genes (*LCIA* and *CAH1*), which showed the same very low expression levels under the three different light intensities used (**Figure 5**). This regulation suggests that unlike the *LHCSR3* genes, the CCM genes may strictly respond to inorganic carbon availability through CIA5-dependent activation. The light effect traditionally ascribed to regulation of the CCM genes (Yamano et al., 2008) may exclusively be associated with changes in intracellular inorganic carbon levels resulting from the changes in the rate of CO_2_ fixation at the different light intensities.

In contrast, *LHCSR1* exhibited very little if any repression in high CO_2_ in both the WT strain and the *cia5* mutant (**Figure 5** and **6**). Slightly lower mRNA accumulation in cells maintained at high CO_2_ could be the consequence of saturating CO_2_ concentrations which promotes fixation, releases electron pressure in the PET chain, and generates less RS. This possibility is supported by the finding that CO_2_ appears to have a stronger effect in ML than in HL (**Figure 5**). Overall, our results suggest that there is an important role for chloroplast-generated signals (i.e., RS) in activation of the photoprotective genes, especially *LHCSR1*, and that this signal may diminish if the cells are exposed to higher levels of CO_2_. Finally, when the cells are transferred from dark to HL, they produce RS in concentrations high enough to reach maximal *LHCSR1* expression in low or high CO_2_; both conditions become limited for CO_2_ fixation which leads to high RS production and maximal activity of *LHCSR1*. Interestingly, *cia5* cells were already able to attain maximal levels of transcript in LL and low CO_2_, potentially a consequence of their reduced ability to concentrate CO_2_ even after the dark pre-acclimation (basal level of the CCM might be higher), which would lead to higher RS in the mutant than in WT cells.

The *PSBS1* gene exhibited some CIA5-dependent induction in low CO_2_ at ML and HL (**Figure 5**). Both *PSBS* genes (*PSBS1*/*2*) contain two conserved enhancer elements (EEC motifs) in their promoter (Correa-Galvis et al., 2016) that are conserved in genes that respond to low CO_2_, such as *LHCSR3* (Maruyama et al., 2014) and various CCM genes (Kucho et al., 2003; Yoshioka et al., 2004). Previous work has suggested that the increases in the levels of the LHCSR3 and PSBS proteins in response to low CO_2_ can be ascribed to those EEC motifs (Correa-Galvis et al., 2016). PSBS protein synthesis is induced in HL, but it is rapidly degraded, except when the cells are incubated under low CO_2_ levels (Correa-Galvis et al., 2016; Tibiletti et al., 2016). The lower induction at LL intensities compared to *LHCSR1* and *LHCSR3* (**Figure 1**), the total lack of repression when PET is blocked by DCMU (**Figure 2**), the strong transcript (**Figure 4**) and protein induction in the presence of UV-B radiation (Allorent et al., 2016), and the regulation of transcript and protein accumulation under low CO_2_ conditions (**Figure 5**), suggest that this protein may be required under extreme conditions when the light intensity is maximal and cells are experiencing photoinhibition.

Together, our data highlight the complex regulatory network that controls expression of the genes encoding proteins that impact NPQ and both the finely tuned and multi-layered regulation that allow cells to acclimate and anticipate HL stress. This regulation is especially relevant in microalgae like Chlamydomonas that are usually exposed to an extremely dynamic light environment. Different Chlamydomonas species can be found in a range of habitats including ponds, lakes, marine waters, wet gardens and agricultural lands, forests, deserts, snow, and even in the air at altitudes of 1100 m (Gorton & Vogelmann, 2003). Therefore, sophisticated regulation for triggering NPQ could allow these organisms to populate various ecosystems, but continued exploration of mechanisms associated with photoacclimation among the varied species is also likely to reveal different suites of strategies to accommodate the absorption of excess excitation energy. Moreover, *LHCSR* and *PSBS* genes are also conserved in mosses like *Physcomitrella patens* (Alboresi et al., 2010) that thrive in environments where the light conditions are very dynamic, while in plants only the PSBS is conserved.

In our work, we have focused on the transcriptional regulation of *LHCSR1*, *LHCSR3*, and *PSBS* genes over the short-term following a dark-to-light transition, concentrating on elucidating how the initial transcriptional activities of these genes are regulated in response to different light intensities, qualities and CO_2_ levels, and how these signaling pathways are linked to photoperception by PHOT1 and UVR8, respond to CO_2_ levels and the role of CIA5 in that response. These analyses indicate that there are multiple pathways that feed into the regulation of the *LHCSR1, LHCSR3* and *PSBS* genes and that there are some clear distinctions in the ways in which expression of these genes are regulated and the regulatory elements involved. The regulatory inputs include photoreceptors, photosynthetic electron flow (e.g., redox, RS), UV-B radiation and CO_2_ levels (the latter, primarily through CIA5). These inputs may be independent, interactive, integrative and compensatory, allowing for optimization of expression with respect to environmental inputs over the course of the day. The factors that regulate the photoprotective genes and the ways in which they are integrated are summarized in the model presented in **Figure 7B**. However, further studies into posttranscriptional regulation of *LHCSR1*, *LHCSR3* and *PSBS* (transcript stability, translation efficiency, protein stability, turnover and modification) under different light and atmospheric conditions over the diel cycle will provide additional critical insights into the integrated regulation that modulates NPQ and protects cells in dynamic light environments.

## MATERIALS AND METHODS

### Chlamydomonas strains

The *Chlamydomonas reinhardtii* strains used in this study were wild-type CC-125 mt^+^ and CC-124 mt^+^ (137c), *phot1* (CC-5392), *uvr8* (CC-5442), *cia5* (CC-2702), the *phot1* rescued strain, designated *pho1t-C*, and the *cia5* rescued strain, designated *cia5-C*. The *phot1* mutant was engineered by CRISPR-CAS9 inactivation (Greiner et al., 2017) and *cia5-C* is described in (Ruiz-Sola et al. 2021). For the complementation of *phot1*, resulting in the *pho1t-C* strain, a 2.25-kb fragment containing the *PHOT1* CDS was amplified by PCR with KOD hot start DNA polymerase (Novagen) using *PHOT1* FW and *PHOT1* RV primers (Table 1), gel purified and cloned into phk330 (Miao et al., 2019) using the BamHI and EcoRI restriction sites for expression under control of *HSP70/RBC* hybrid promoter. Junctions and inserts were sequenced, and constructs were linearized by KpnI before transformation into the *phot1* mutant. Eleven ng/kb of linearized plasmid (Mackinder et al., 2016) was mixed with 400 μL of 1.0 x10^7^ cells mL^-1^ and electroporated in a volume of 120 µL in a 2-mm-gap electro-cuvette using a NEPA21 square-pulse electroporator (NEPAGENE, Japan). The electroporation parameters were set as follows: Poring Pulse (300 V; 8 ms length; 50 ms interval; one pulse; 40% decay rate; + Polarity), Transfer Pulse (20 V; 50 ms length; 50 ms interval; five pulses; 40% decay rate; +/-Polarity). Transformants were selected on solid agar plates containing 7.5 μg/ml zeocin and screened based on their NPQ capacity using the following protocol: transformants grown in liquid TAP medium for three days in 96-well transparent microplates were shifted to high-salt medium (HSM) medium and exposed to 300 μmol photons m^-2^ s^-1^ for 4 h before measuring NPQ using a Maxi-Imaging PAM fluorometer (see Chlorophyll fluorescence analysis paragraph below). Colonies with WT-levels of NPQ were chosen as putative complemented strains. This was further confirmed by western blot analyses using anti-PHOT antiserum (LOV1 domain) as previously described (Zorin et al., 2009).

**Table 1.**
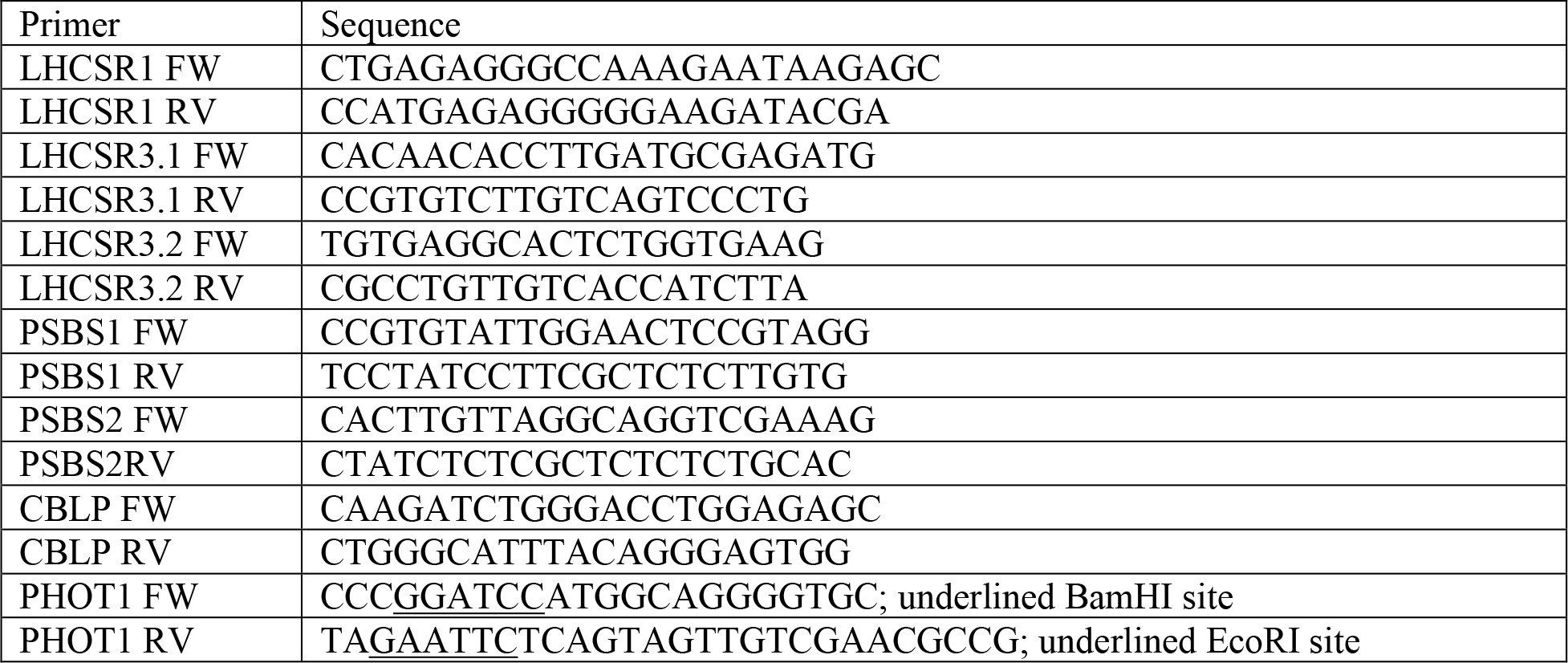
Primer sequences used for transcript quantifications.

### Growth conditions and induction treatments

Cells were grown to mid-exponential phase (∼10 µg/ml chl) at 23°C under continuous white LED light (30 µmol photons m^-2^ s^-1^) with shaking at 130 rpm in 50 ml TAP medium (Harris 2001) in 250 ml flasks. The spectra of the light sources in the growth chambers are shown in **Supplementary Figure 7**, with the spectrum of sunlight shown for comparison. Prior to experimental treatments, the cells were adjusted in TAP medium to a chl concentration of 10 µg ml^-1^ and acclimated in the dark for 24 h to lower the levels of the *LHCSR1*, *LHCSR3.1, LHCSR3.2, PSBS1* and *PBS2* transcripts. Cells were then harvested by centrifugation (3000 × g, 1.5 min) at 23°C, washed once at room temperature with minimum medium (high-salt medium, HSM) and then resuspended in HSM and kept shaking for two additional hours in the dark (to further lower transcript levels). After various treatments, described in the ‘RESULTS’ section, the cells were harvested by centrifugation (3000 × g, 1.5 min), flash frozen with liquid nitrogen and stored at −80°C until the RNA was extracted. UV-B radiation at a level present in natural sunlight at midday (200 µW/cm^2^) was from a Philips TL20W/01RS narrowband UV-B tube with half maximal transmission at 311 nm. Control samples were maintained under a UV-B protective plexiglass filter. Experiments using low light (LL, 30 µmol photons m^-2^ s^-1^), high light (HL, 480 µmol photons m^-2^ s^-1^), very high light (VHL, 960 or 1,000 µmol photons m^-2^ s^-1^) and stepped light levels (from 5-960 µmol photons m^-2^ s^-1^) were as described in the text, while exposure to blue (450 nm peak) and red (660 nm peak) light (**Supplementary Figure 3**) were in a HiPoint plant growth chamber (FH-1200). To suppress photosynthetic electron flow, DCMU was added to cultures (to 10 µM) in the dark immediately prior to placing them under the various conditions of illumination.

### Evaluating photosynthetically active and UV-B radiation over the diel cycle

PAR and UV-B intensities were measured during one week in July in California, from sunrise to sunset, using a LI-250A Light Meter (LI-COR®) and an Inc Solarmeter Model 6.2 (Solar Light Company), respectively. The curves in **Figure 1A** show the intensity (in µmol photons m^-2^ s^-1^, also designated µE) of photosynthetically active radiation (PAR) and the UV-B radiation (power density in µW/cm^2^) over the course of a representative day from 6:00 AM to 8:00 PM.

### RNA extraction and quantitative RT-PCR

Total RNA was isolated using a phenol/chloroform based protocol (Sanz-Luque & Montaigu, 2018). Residual DNA was removed by TURBO^TM^ DNase (Thermo Fisher Scientific) and cDNA synthesized by reverse transcription of 1 µg of isolated total RNA using the iScript^TM^ Reverse Transcription Supermix (Bio-Rad) in a 20 µl reaction volume. cDNA was diluted by a factor of 2.5 and then 1 µl of the resulting 50 µl (a total of ∼20 ng cDNA) served as the template in a 20 µl RT-PCR reaction. Real-time PCR was performed with the SensiFast^TM^ SYBR No-Rox Kit (Bioline) in a Roche Light Cycler 480 as described by the manufacturer. A 2-step cycling condition was used (2 min 95°C, 40 cycles of 95°C 5 s, 60°C 30 s) with the fluorescence yield quantified at the end of each cycle. The *CBLP* gene served as the housekeeping control and relative fold differences were calculated based on the ΔCt method (2^-(Ct target gene - Ct *CBPL)*^) (Livak & Schmittgen, 2001; Schmittgen & Livak, 2008). The primer sequences for transcript quantification are displayed in **Table 1**; specific primer pairs were used to distinguish *LHCSR3.1* and *LHCSR3.2* transcripts and *PSBS1* and *PSBS2* transcripts.

### Statistics

Statistical analysis of the data was performed with GraphPad PRISM8 software (8.4.1) with one- or two-way ANOVA using Tukey’s post-hoc test or Uncorrected Fisher’s LSD. The significance of differences between treatments are given as ANOVA-derived p-values that are depicted in the figures as *, **, or ***, representing 0.05, 0.005 and 0.001, respectively. p-values are also indicated in the text.

## SUPPLEMENTARY DATA

**Supplementary Figure 1:**
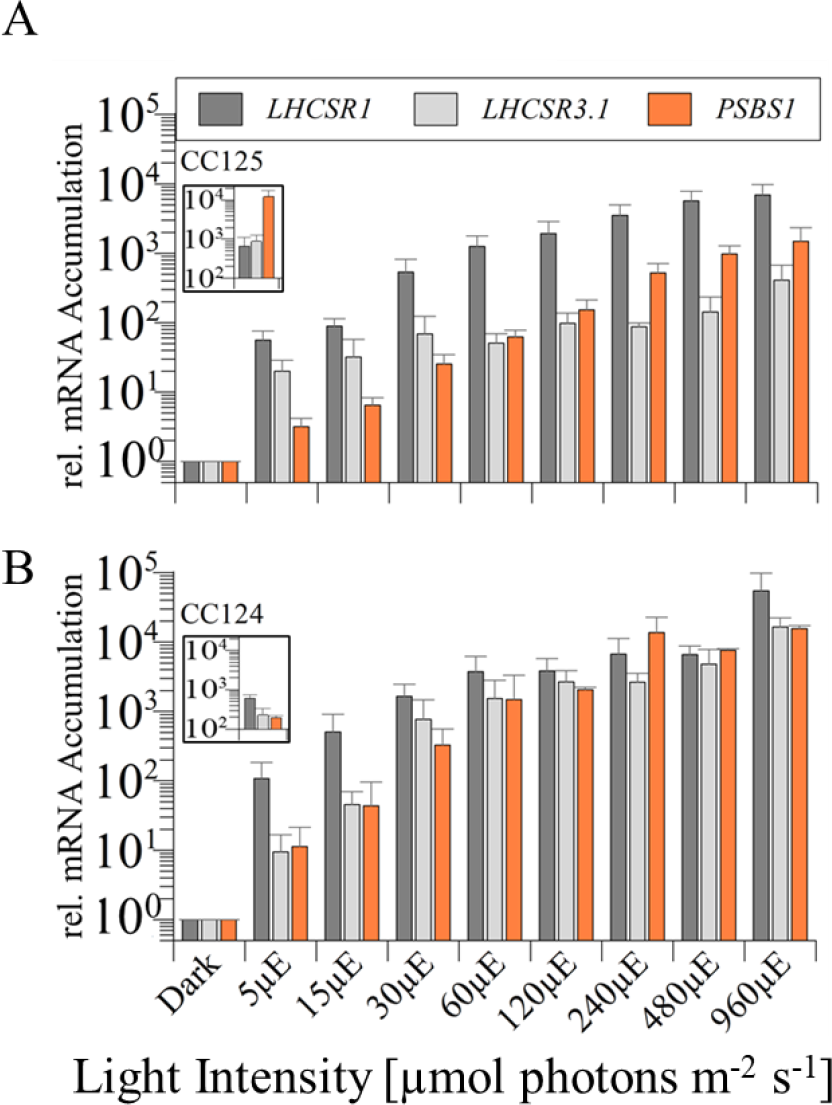
Changes in levels of *LHCSR* and *PSBS* transcripts in CC125 and CC124 after one hour of irradiation at indicated light intensities. *LHCSR1*, *LHCSR3.1* and *PSBS1* transcripts levels were quantified following a 1 h incubation at different light intensities. The **A**) CC-125 (replotted from **Figure 1**) and **B**) CC-124 strains were grown in TAP at LL (30 µmol photons m^-2^ s^-1^) and then transferred for 24 h to TAP in the dark. Following this incubation, the cells were transferred to HSM (photoautotrophic conditions) and kept for two additional hours in the dark prior to a 1-h light exposure at each of the indicated intensities. Values for each of the three transcripts were normalized to the dark value. The insets in each panel show the initial levels of transcripts determined for cells in the dark before the 1 h treatment period, without normalizing the data. n=3+SD. Statistical analyses and P-values are listed in **Supplementary Table S1.**

**Supplementary Figure 2:**
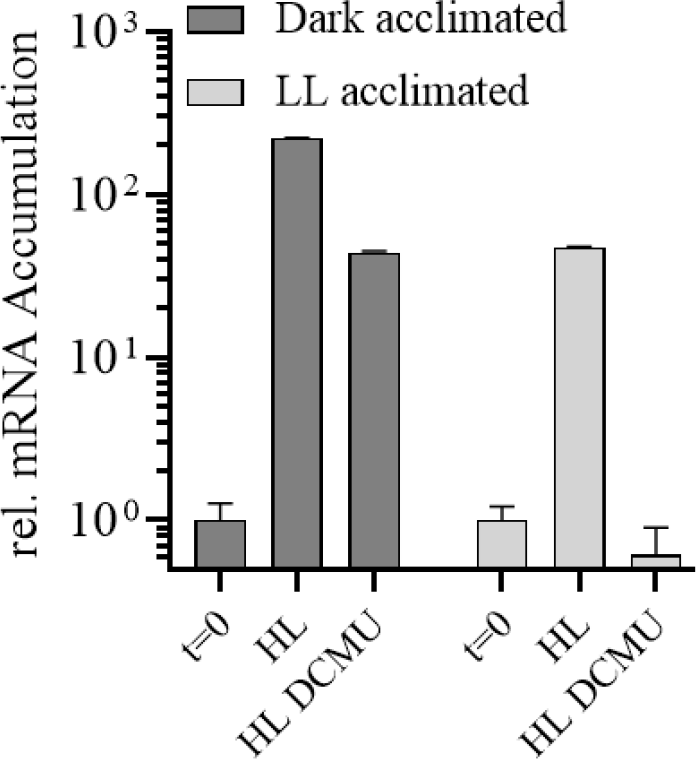
Induction of *LHCSR3.1* transcript. CC-125 cells were acclimated in HSM overnight in darkness or LL (15 µmol photons m^-2^ s^-1^). The next morning the samples were taken for t=0 and cells were exposed to HL (300 µmol photons m^-2^ s^-1^) for 1 h in the presence or absence of 40 µM DCMU. n=3+SD. Statistical analyses and P-values are listed in **Supplementary Table S2.**

**Supplementary Figure 3:**
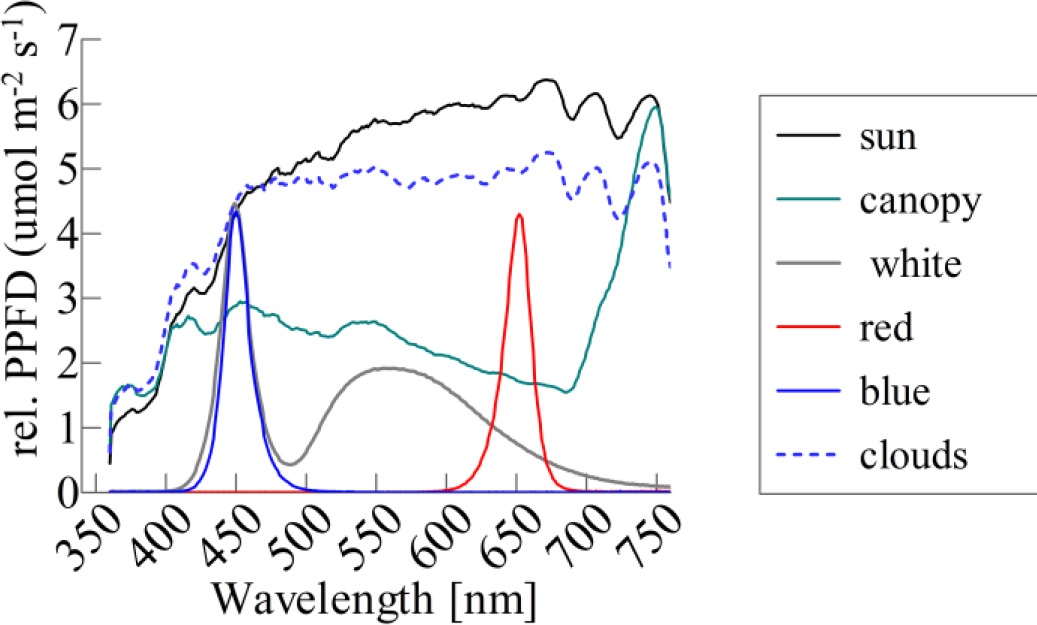
Spectra of light in the environment and the light used for growth in the chambers. Spectra of white, red, and blue LEDs used for cell treatment in the growth chambers compared to the spectrum for sunlight, which was measured during the afternoon in full sun and in shade beneath a tree (5:00 PM, California, 2020). Cells were grown in white LED light (grey line). The white LED within the growth chamber is 6000 K, which is highly similar to sunlight/daylight LEDs (6500 K), although the sunlight spectrum is more evenly distributed over the PAR spectrum (400-760 nm).

**Supplementary Figure 4:**
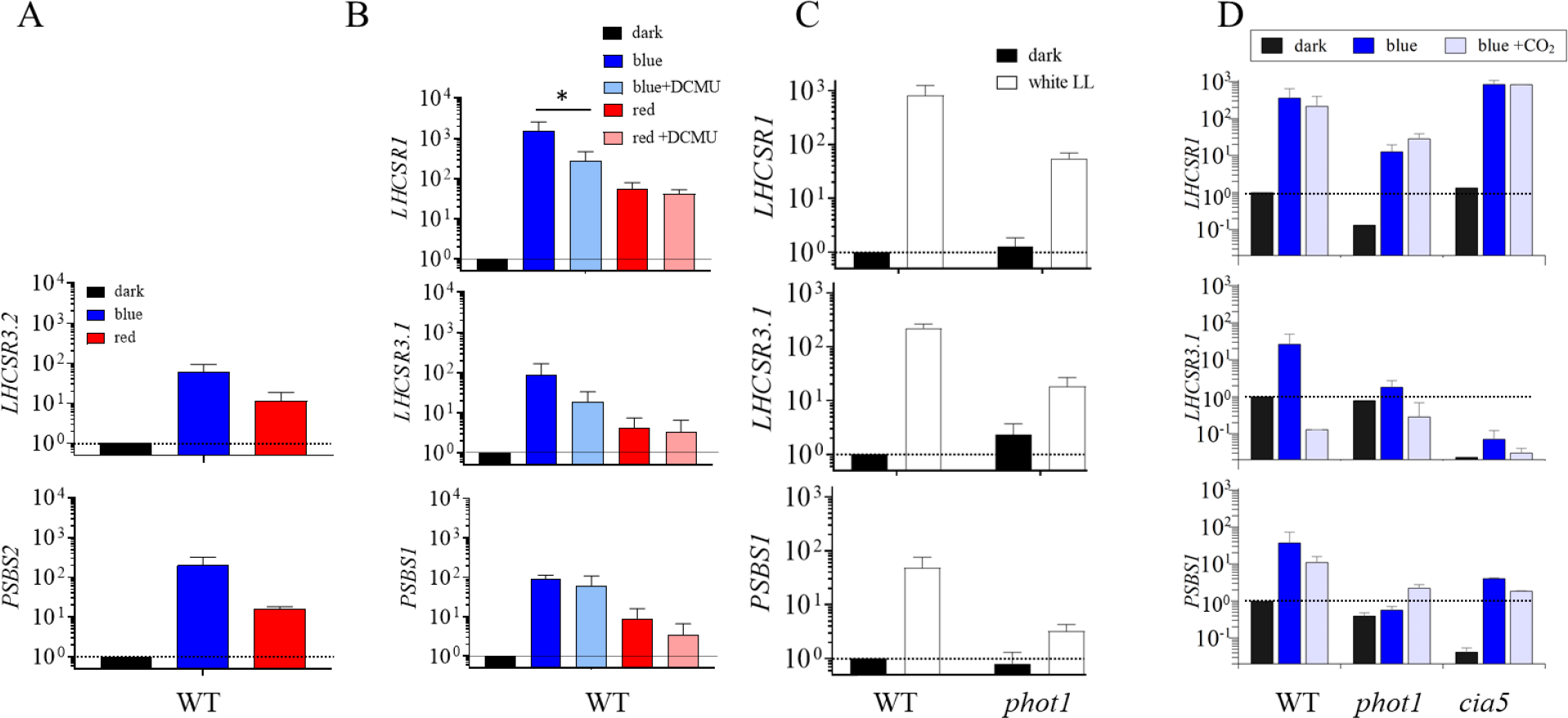
The effects of light quality, electron flow and CO2 on accumulation of transcripts encoded by the photoprotective genes. A) Induction of *LHCSR3.2* and *PSBS2* genes following 1 h of blue and red light. B) Induction of the *LHCSR1*, *LHCSR3.1* and *PSBS1*genes after 1 h of blue or red light in the presence or absence of DCMU (10 mM). C) Induction of *LHCSR1*, *LHCSR3.1* and *PSBS1* genes in WT and the *phot1* mutant for 1 h in LL. D) Induction of *LHCSR1*, *LHCSR3.1* and *PSBS1* genes in WT and the *phot1* and *cia5* mutants for 1 h in blue light in the presence or absence of 5% CO_2_. CC-125 WT, *phot1* and *cia5* cells were grown as described in the legend of Figure 1 before the various treatments. The light intensities were 30 µmol photons m^-2^ s^-1^ white (white bar), blue (blue bar) or red (red bar) light. n=3+SD. Statistical analyses and P-values are listed in Supplementary **Table S4**.

**Supplementary Figure 5:**
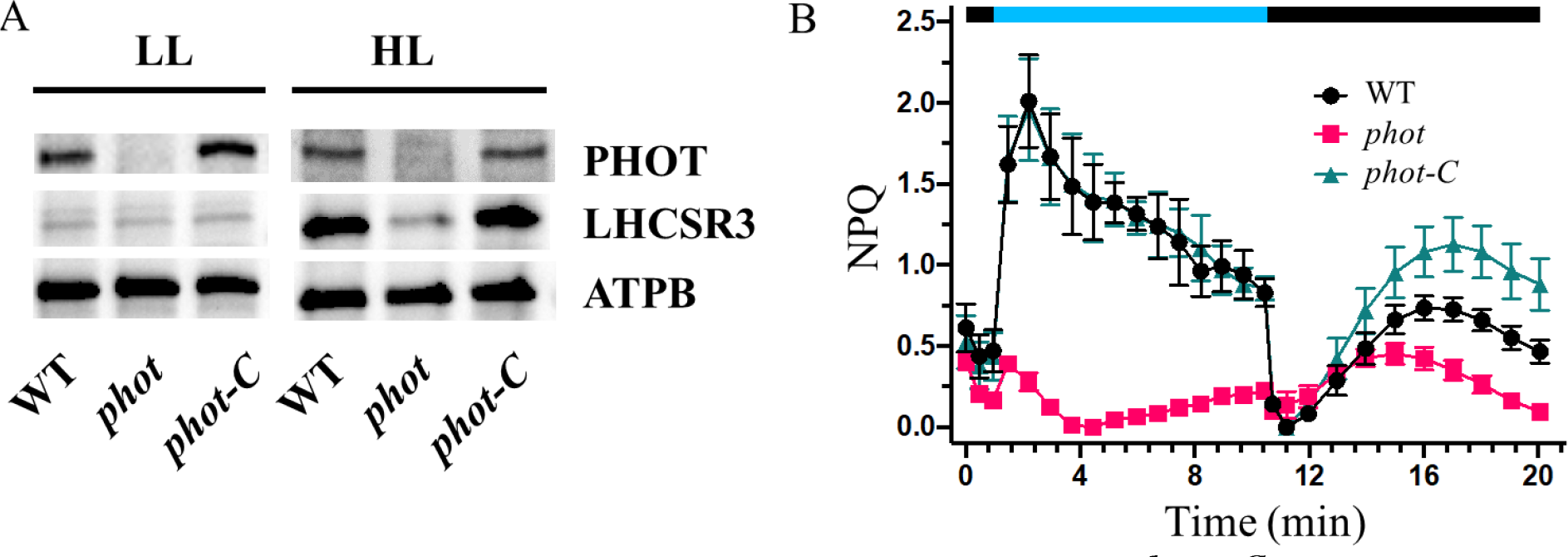
Phenotypes of new *phot1* mutant and the *phot1-C* complemented strain. **A)** Immunoblot blot analyses of WT, *phot1* and *phot1-C* acclimated to LL (15 µmol photons m^-2^ s^-1^) and after exposure to HL (300 µmol photons m^-2^ s^-1^) for 4 h; the proteins examined were PHOT, ATPB and LHCSR3; ATPB served as loading control. **B**) Kinetics of NPQ for WT, *phot* and *phot1-C* strains after exposure to HL for 4 h. NPQ was recorded over 10 min of illumination at 600 µmol photons m^-2^ s^-1^ of blue light (blue bar), followed by 10 min of darkness (black bar), during which relaxation of NPQ was monitored. Values plotted are the means of nine replicates +/- SD (three biological replicates each one measured at three technical replicates). Before the NPQ measurements the cells were shaken in the dark for 20 min. The phenotypes of these strains are in accord with what has been previously reported for similar strains (Petroutsos et al., 2016).

**Supplementary Figure 6:**
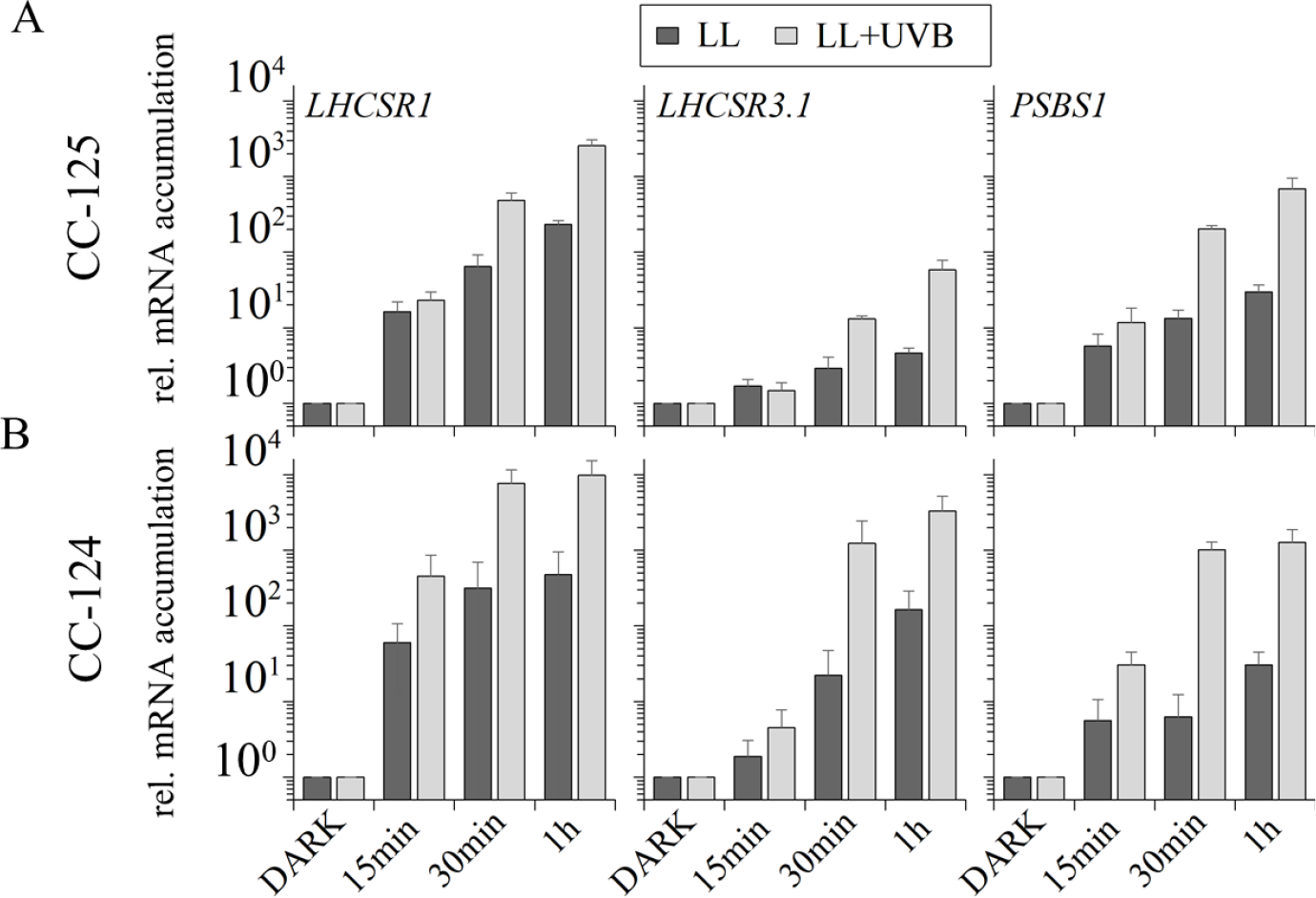
Induction of *LHCSR* and *PSBS* genes at various times in LL in the presence or absence of UV-B irradiation. **A**) WT CC-125 or **B**) WT CC-124 cells were grown as described in the legend of **Figure 1** prior to exposure to LL (30 µmol photons m^- 2^ s^-1^) or LL+UV-B (30 µmol photons m^-2^ s^-1^ PAR plus 200 µW/cm^2^ UV-B) for up to 1 h. Statistical analyses and P-values are listed in **Supplementary Table S6.**

**Supplementary Figure 7:**
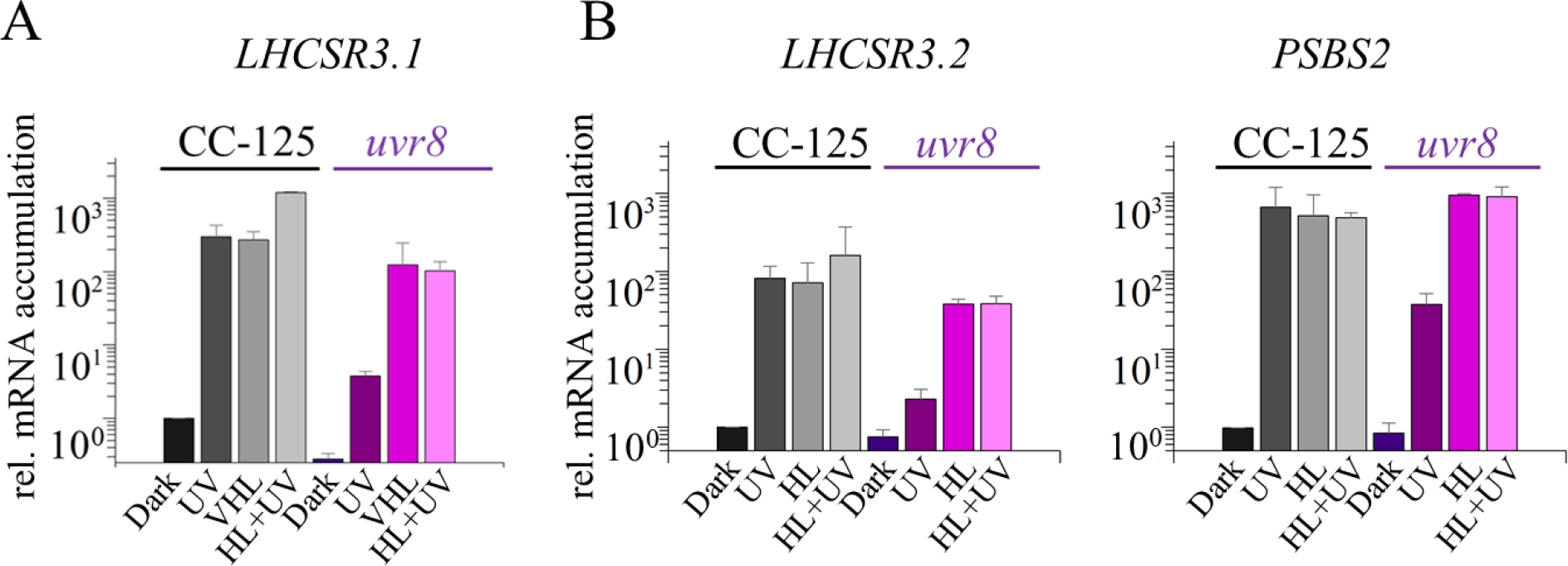
Impact of UV-B radiation on expression of photoprotective genes. **A**) Changes in levels of *LHCSR3.1* transcript 1 h after transfer to darkness (Dark), UV-B (UV; 200 µW/cm^2^) radiation, VHL (1000 µmol photons m^-2^ s^-1^), or VHL + UV-B radiation. **B**) *LHCSR3.2* and *PSBS2* after 1 h of darkness, UV-B (UV) radiation, HL (480 µmol photons m^-2^ s^-1^), or HL+UV-B radiation. WT CC-125 (grey-black bars) and *uvr8* cells (coloured bars) were grown as described in the legend of **Figure 1**. Cultures were grown as described in the legend of Figure 1, divided and either kept in the dark or subjected to the various light treatments described. n=3+SD. Statistical analyses and P-values are listed in **Supplementary Table S7.**

## SUPPLEMENTARY METHODS

### Protein analysis

Protein samples of whole cell extracts (0.5 µg chl) were loaded on Mini-PROTEAN TGX, 4- 20% Biorad precast gels and blotted onto nitrocellulose membranes. Antisera against LHCSR3 and ATPB were from Agrisera (Vännäs, Sweden) while the PHOT1 antiserum (LOV1 domain) was previously described (Zorin et al., 2009). ATPB was used as a loading control. The secondary anti-rabbit antibodies were conjugated to horseradish peroxidase. Immunoblots were developed with the ECL detection reagent and images were obtained using a CCD imager (ChemiDoc MP System, Bio-Rad).

### Chlorophyll fluorescence analysis

Fluorescence-based measurements of photosynthetic parameters were performed with a Maxi-Imaging PAM fluorometer (Heinz Walz GmbH, Effeltrich, Germany). NPQ was calculated using the equation (Fm-Fm’)/Fm’. Fm and Fm’ are maximum fluorescence yields after dark preincubation and in steady state light, respectively, measured using a saturating light pulse. Before assaying NPQ, the cells were exposed to high intensity actinic light (600 µmol photons m^-2^ s^-1^) for 4 h to induce *LHCSR3* and then dark acclimated with vigorous shaking for 20 min.

## Funding

All authors thank the Human Frontiers Science Program RGP0046/2018 for funding this work. E.S.-L. thanks the European Union’s Horizon 2020 research and innovation program under the Marie Sklodowska-Curie grant agreement no. 751039 and the ‘Plan Propio UCO’ program from University of Cordoba, Spain for postdoctoral support. Support was also provided to A.R.G. by the Carnegie Institution for Science. D.P., G.V. and Y.Y. would also like to thank for funding the French National Research Agency (i) in the framework of the Young Investigators program ANR-18-CE20-0006 through the funding of the project MetaboLight, (ii) in the framework of the Investissements d’Avenir program ANR-15-IDEX-02, through the funding of the “Origin of Life” project of the Univ. Grenoble-Alpes and (iii) through the funding of the Grenoble Alliance for Integrated Structural & Cell Biology (GRAL) project ANR-17-EURE-0003.

## Author contributions

Conceptualization: P.R., E.S.-L., D.P. and A.R.G.

Methodology: P.R., E.S.-L., D.P.

Investigation: P.R., E.S.-L., Y.Y and G.V.

Supervision: D.P., A.R.G.

Writing—original draft: P.R., E.S.L., A.R.G.

Writing—review & editing: P.R., E.S.-L., D.P., A.R.G.

## Competing interests

Authors declare that they have no competing interests.

## Notes

### Competing Interest Statement

The authors have declared no competing interest.

### Summary of Updates

misspelling in Title

